# MotSASi: Functional Short Linear Motifs (SLiMs) prediction based on genomic single nucleotide variants and structural data

**DOI:** 10.1101/2021.08.05.455287

**Authors:** Mariano Martín, Carlos P. Modenutti, Juan P. Nicola, Marcelo A. Marti

**Author notes:** Mariano Martín,; Marcelo A Marti.

## Abstract

Short linear motifs (SLiMs) are key to cell physiology mediating reversible protein-protein interactions. Precise identification of SLiMs remains a challenge, being the main drawback of most bioinformatic prediction tools their low specificity (high number of false positives). An important, usually overlooked, aspect is the relation between SLiMs mutations and disease. The presence of variants in each residue position can be used to assess the relevance of the corresponding residue(s) for protein function, and its (in)tolerance to change.

In the present work, we combined sequence variant information and structural analysis of the energetic impact of single amino acid substitution (SAS) in SLiM-Receptor complex structure, and showed that it significantly improves prediction of true functional SLiMs. Our strategy is based on building a SAS tolerance matrix that shows, for each position, whether one of the possible 19 SAS is tolerated or not. Herein we present the MotSASi strategy and analyze in detail 4 SLiMs involved in intracellular protein trafficking. Our results show that inclusion of variant and sequence information significantly improves both prediction of true SLiMs and rejection of false positives, while also allowing better classification of variants inside SLiMs, a results with a direct impact in clinical genomics.

## Introduction

Short linear motifs (SLiMs) mediate different processes that are key to cell physiology. These short stretches of amino acid sequences are frequently located within intrinsically disordered regions of proteins, or in exposed flexible loops, that allow them to interact with their binding partners. These transient and reversible SLiM-mediated protein-protein interactions (PPIs) are central for controlling protein stability and subcellular localization, as well as for building the dynamic protein complexes that regulate multiple cellular processes. Post-translational modification is also controlled by SLiMs, allowing corresponding enzymes (phosphatases, kinases, etc.) to recognize their substrates, and by these means, modulate and integrate different signals that are transmitted to the wider protein population in order to regulate the cell decision-making. Intracellular protein trafficking is also directly regulated by SLiMs, since the presence (or absence) of certain SLiMs determines the subcellular localization of proteins, thereby organizing and determining organelle function (Van Roey et al., 2014). Clearly, the relevance and ubiquity of SLiMs make their identification and characterization a very important endeavor.

Precise identification of SLiMs remains a major challenge. High-throughput screening experiments and computational studies estimate that the human proteome is predicted to contain over a million of binding motifs (Tompa et al., 2014). However, the relevance of each of these predicted motifs for the cell physiology is difficult to address without experimental confirmation. SLiMs have been historically characterized using cell biology and biophysical approaches, and is limited by our inability to characterize SLiMs in *in vivo* models in the context of complex multiprotein assemblies, and the inherent difficulty of reproducing these assemblies in vitro. Recently, we identified and functionally characterized three different SLiMs involved in the intracellular trafficking and membrane expression of the sodium/iodide symporter (NIS) (Martín et al., 2019, 2021a, 2021b), a key molecule in thyroid pathophysiology.

Initial SLiMs prediction is most commonly addressed by bioinformatic tools. There are at least eight softwares and forty packages for the prediction of new SLiMs (Edwards and Palopoli, 2015). All of them scan a given query sequence for matches against a database of characterized motifs, usually defined by a regular expression. Additionally, most of them use residue conservation scores and sequence secondary structure analysis for improving their predictions. The Eukaryotic Linear Motif (ELM) resource has one of the biggest SLiMs databases and one of the most used motif prediction software (Kumar et al., 2019). It uses logical filters, such as taxonomic range, protein structure, subcellular localization, presence of globular domains and sequence probability, in order to improve the precision of each motif prediction. Another relevant tool is SLiMSuite, a toolkit for Short Linear Motif discovery and molecular evolution that presents different packages for motif prediction (Edwards and Palopoli, 2015). The toolkit makes use of evolutionary relationships, statistical models, protein features and machine learning algorithms for making better SLiMs prediction. Another widely used prediction tool is MiniMotif Miner (MnM) (Lyon et al., 2018). This tool uses different filters that improve accuracy: proteome frequency, protein surface prediction, evolutionary conservation, protein–protein interactions, related molecular functions, genetic interactions and secondary structure filters, reaching an accuracy of about 90%. Furthermore, this software evaluates the presence of SNPs that disrupt or generate motifs but does not consider them for the motif prediction. Although this is an interesting add up, the tool considers that all SNPs affecting a motif always eliminates it without considering that many missense mutations do not affect the motif because they take place in flexible positions, or the change is tolerated by the robustness of the motif-domain interaction. The MEME suite and PSSMSearch are other frequently used software for SLiM discovery (Bailey et al., 2015; Krystkowiak et al., 2018). These tools use Position-Specific Scoring Matrices (also called Profiles or Position Weight Matrices) trained on input sequences for novel motif prediction. The D-MIST method also uses profiles but is based on motif-domain structural topology. This technique extracts motifs from structural data and converts them in sequence profile in the form of PSSMs. The PSSMs can be improved with a Gibbs sampling in the set of proteins that interact with proteins that contain the domain of interest. D-MIST has been only used for sampling yeast PPIs (Betel et al., 2007). Other common tools used for motif search and discovery are ScanProsite and DILIMOT (Sigrist, 2002; Neduva and Russell, 2006).

The main drawback of most SLiMs prediction tools is their low specificity, due to the high number of false positives. Therefore, predictions are usually taken as working hypotheses that move forward towards experimental validation. Reasons underlying false positive predictions are the large (or huge) number of available sequences (i.e search space), the degenerate nature of the short motif, and most important the fact that for many SLiMs, the number of experimentally validated, true positives, SLiMs is rather small, limiting a precise definition of the motif. As previously mentioned, predictions are usually improved using additional filters. In this context, in the present work, we show that human SLiMs prediction can be improved by developing and applying filters related to: i) the motif structure in the context of its interaction with the corresponding protein receptor, and ii) its relation to the cause of genetic based diseases.

Given their critical roles in cellular physiology, mutations disrupting SLiMs function lead to several human diseases (Deretic et al., 1998; Müller et al., 2003; Kalay et al., 2005; Etxebarria et al., 2015; Martín et al., 2021a). Where the SLiM is essential for the corresponding protein function, motif disruption results in a non-functional protein and the observed phenotype is similar to that observed for null mutations. Such is the case of mutation in the tyrosine-based motif of the Low-density lipoprotein receptor (LDLR), where disruption of the motif conducts to a deficient receptor internalization, leading to familial hypercholesterolemia (Etxebarria et al., 2015). In other cases, motif disruption alters only some particular aspect of the corresponding protein function, such as protein stability or subcellular localization, and therefore a specific motif disruption-associated phenotype is observed. An example is the atypical Usher Syndrome caused by disruption of Usher Syndrome type-1G protein (USHG1) PDZ-binding motif and therefore, the interaction with Harmonin (USH1C), leading to a deficient assembly of the protein network that mediates mechanotransduction in cochlear hair cells (Kalay et al., 2005).

The relation between SLiMs mutations and disease, as will be shown in the present work, can be used for two key developments, which indeed are two faces of the same coin. On one hand, in the context of next generation based human sequencing for molecular diagnosis, it is critical to determine whether a variant (mutation) found in a given patient is pathogenic (i.e leads to disease development), or benign (has no effect). Although clinical criteria for pathogenicity includes other properties (family segregation, relationship between observed and expected phenotype, etc.) from a molecular viewpoint, they result from a significant alteration of the corresponding protein function. Prediction of whether a particular variant affects protein function is not straight forward for those resulting in Single Amino Acid Substitutions (SAS), and there are many bioinformatic tools that perform this task (Adzhubei et al., 2010; Vaser et al., 2016). Most of them are based on residue conservation and may include additional features, some of which are related to protein structure. However, to our knowledge, explicit inclusion of SLiMs, and their properties, as presently proposed, had not been performed. On the other hand, the presence of either benign or pathogenic variants in each residue position can be used to assess the relevance of the corresponding residue(s) for protein function, and its (in)tolerance to change. Therefore, we hypothesized that there should be a differential pattern of pathogenic/benign variations in true SLiMs with respect to those sequences that are identified as SLiMs by sequence analysis, but are not functional (i.e false positives).

In the present work, we show that combined use of sequence nucleotide variant information, such as Clinical Significance and allele frequency, with structural-based analysis of the energetic impact of each SAS in the corresponding SLiM-Receptor complex structure improves SLiMs prediction confidence (i.e identification of true functional motifs). Our strategy is based on building a SAS tolerance matrix that shows for each SLiM sequence position whether one of the possible 19 SAS is tolerated or not. Furthermore, these matrices allow a deeper understanding of each analyzed SLiM possible sequence, as well as providing support to novel variant classification in a clinical context. Herein, we present our MotSASi (**<u>Mot</u>**if-occurring **<u>S</u>**ingle **<u>A</u>**mino acid **<u>S</u>**ubstitution **i**nformation) method and thoroughly analyze 4 SliMs involved in intracellular trafficking: tyrosine based motif (NPx[Y/F]), type 1 PDZ-binding motif ([S/T]x[V/I/L]COOH), tryptophan-acidic motif ([L/M]xW[D/E]) and acidic di-leucine motif ([E/D/Q/R]xxxL[L/I/V]). Our method allowed the identification of a significant number of novel motifs with high confidence, expanding the original true positive set of motif-containing proteins from 2 to 10 times. Moreover, in most of the cases, we discarded 10-20% of the initially predicted motifs in the human proteome. The implementation of our MotSASi method will significantly improve the prediction of biologically relevant motifs, thus avoiding the experimental study of non-relevant motifs.

## Results

The results are organized as follows: First, we present the general aspects of our SLiMs characterization pipeline, and analyze how the number of highly confident predicted SLiMs, as well as the number of false positives evolve for all analyzed motifs. Second, using the phospho-independent tyrosine-based motif, we show how SAS pathogenic-benign matrix evolves as additional data and particular motifs are added. Finally, we show and discuss in detail the results (newly identified proteins with high confidence SLiMs and the SAS pathogenicity matrix) for all the analyzed motifs.

### SLiM characterization pipeline

The process begins with a sequence based query of each motif in the whole human proteome, consisting of all protein sequences annotated in the *Homo sapiens* genome in the UniProt database, using regular expressions (for each motif regular expression see methods section) (Fig 1 and Table 1). Identified putative motifs correspond to our Initial Set. Those motifs that present at least one variant with Clinical Significance (ClinSig) or high Allelic Frequency (AF), constitute our universe of analyzable motifs and were named Set 0 (S0). Subsequently, we analyzed whether experimentally validated motifs (as reported in ELM database or bibliography) were identified by our search and assigned them as Positive Set 0 (P0). As shown in Table 1, currently less than 3% of the potential motifs were experimentally validated, underscoring the relevance of better identification tools. P0 corresponds to our initial positive control group, and was used to determine: i) the degree of conservation, ii) the secondary structure and iii) the position of the motif along the protein. Furthermore, using data from ClinVar and GnomAD, we constructed two initial SAS matrices. One, with the clinical significance of the variant [Pathogenic (P) or Benign (B)] which we will refer to as ClinSig matrix, and the second with the allele frequency (represented as −log10(AF)), which we will refer to as AF matrix. As expected, initial matrices contained several missing values (see S3, S5 and S7 Figs). Usually, less than 3% of all potential motifs have SAS with ClinVar entry and, although 80-90% display frequency values in GenomAD, only 10% have high AF and can thus be considered as tolerated (see below).

**Table 1.** Summary of the results obtained with the MotSASi method.

| Motif Name | Tyrosine based | Type 1 PDZ-binding | Tryptophan acidic | Acidic di-Leucine |
| --- | --- | --- | --- | --- |
| Consensus Sequence | NPx[Y/F] | [S/T]x[V/I/L] | [L/M]xW[D/E] | [E/D/Q/R]xxxL[L/I/V] |
| Initial Set | 1823 | 1022 | 1894 | 47821 |
| S0 | 161 | 81 | 114 | 6675 |
| **P0** | 17 | 71 | 8 | 20 |
| **N1f** | 25 | 5 | 1 | 0 |
| **P1f** | 17 | 75 | 8 | 20 |
| **S1f** | 135 | 56 | 112 | 6672 |
| **N2f** | 88 | 14 | 54 | 2092 |
| **P2f** | 18 | 86 | 8 | 83 |
| **S2f** | 71 | 36 | 59 | 4517 |
| **N3f** | 147 | 22 | 103 | 6297 |
| **P3f** | 28 | 96 | 18 | 171 |
| **S3f** | 2 | 18 | 0 | 204 |
| **Nf †** | 341 | 22 | 201 | 8522 |
| **Pf** | 28 | 96 | 18 | 171 |
| **Remaining Set ‡** | 1464 | 975 | 1675 | 39299 |

† Discarded with the use of the final matrix due to incoherent variants and the final definition of the consensus sequence. ‡ Protein that remains to be analyzed from the Initial Set due to a lack of reported variants.

**Figure 1.**
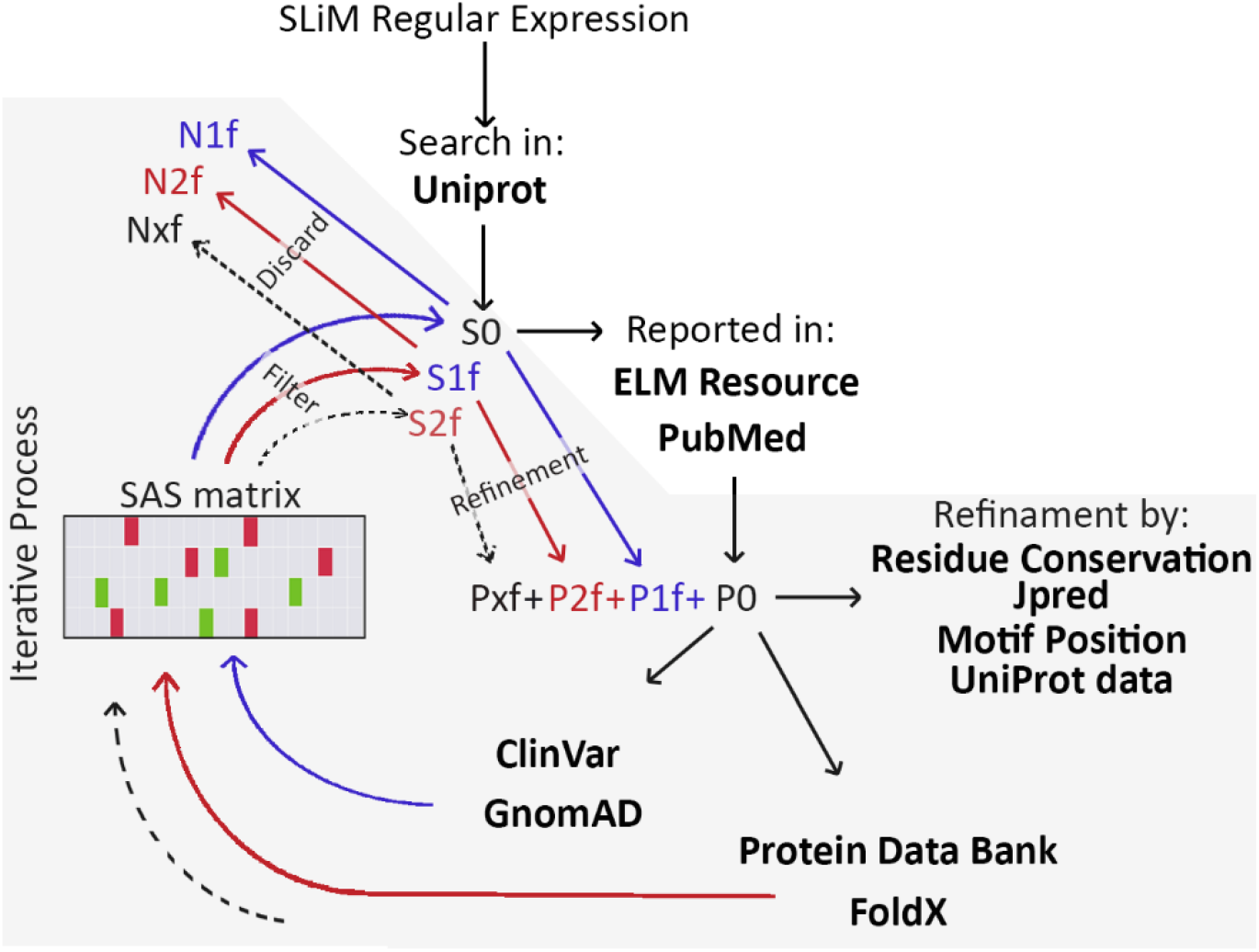
MotSASi’s Pipeline. Each SLiM regular expression is searched independently in the human proteome. Positive datasets (Px) variants information (ClinVar GenomAd), motif containing protein characteristics (Jpred, Uniprot), are used to iteratively build the SAS ClinSig, AF and ΔΔG (using PDB and FoldX) matrices, classify all potential motifs, build the final SAS tolerance matrix and final Positive (Pf), Discarded (Df) and remaining sets (Sf).

Allele frequency is a known proxy for clinical significance of a variant. According to the American College of Medical Genetics and Genomics (ACMG) criteria, an AF over 5% is enough to classify a variant as bening, while AF higher than those expected for a recessive disorder, or presence of homozygosity for dominant cases, are strong criteria towards a benign classification (Richards et al., 2015). The opposite case is more complex, while pathogenic variants should have low AF, there are also many benign variants which just have low frequencies out of chance. To use this information in the context of SLiMs, we assumed that SAS showing high AF in the P0, evidence that this SAS is tolerated, while those SAS assigned as P in ClinVar and showing only low AF values in the PS1 set are not. We define a tolerated SAS if it is shown to occur in the context of the SLIMs but does not affect its function significantly (i.e., a proxy for the underlying variants as being benign). To define a AF cut-off value, we analyzed the −log10(AF) distribution (for visual and analyzable purpose we used the −log10(AF), where low values of AF are represented as higher number in −log10(AF) and vice versa) of pathogenic and benign variants in the whole human proteome (S1 Fig). Our data shows that −log10(AF) of 4 is a good cut-off value. Therefore, we considered a variant with a −log10(AF) < 4 as benign, while a value > 4 was considered as either pathogenic or benign. With this information, we analyzed all S0 motif sequences (except those already in P0) and classified them as either: Negative 1 (N1), S1 and P1

In this first filter, we dismissed the variants occurring at the flexible positions inside the motifs (the x in the motif consensus sequence). N1 corresponds to those motifs from S0 which show one (or more) variants assigned as bening (either directly from ClinVar and/or from AF analysis) in a non tolerated position (in the SAS matrix), or as pathogenic (directly from ClinVar) in a tolerated position. P1 corresponds to those potentially identified motifs which show the presence of variants for which there is information in the SAS matrices, showing no contradiction in their classification. In this case, the higher the number of known variants in the potential motif whose characteristics agree with those of the P0, the higher the confidence as a true SLiM. Finally, S1 are those remaining motifs for which no comparison could be performed.

Set P1 was further refined using the motif conservation score, secondary structure, relative position in the protein, and known protein localization filters. Those motifs from P1 whose characteristics are not in accordance with those of experimentally tested motifs were moved to N1. After filtering, the variants of the new predicted motifs (P1) were added to the ClinVar and AF SAS matrices, and a new classification round was performed. We continued with this iterative process until no changes were observed. These final, refined, sets are referred to as N1f, S1f and P1f.

As already mentioned, SLiMs work by physically interacting with their protein receptors, and thus SAS that hinder this interaction render the motif non functional. For many motifs, complex structures between the receptor and either a small peptide including the SLiM, or the whole protein harboring it, are available. To evaluate how the SAS in the motif affects protein-protein interaction, we computed the change in the binding free energy (ΔΔG) between wild type (or reference) motif sequence and each possible SAS using the FoldX software. As shown in several works (Schymkowitz et al., 2005; Radusky et al., 2018), a large positive ΔΔG suggest that SAS results in a disruption of the corresponding protein-protein interaction, thus indicating a potential pathogenic variant, while values close to zero suggest the change is well tolerated. Thereafter, using structural information, we built for each motif and all possible SAS the corresponding ΔΔG matrix (see S3, S5 and S7 Figs), and used it to further classify the motifs in S1f and P1f.

In a first step, we considered SAS matrix positions as non tolerant where ΔΔG is higher than 2 kcal/mol and as tolerant those positions with ΔΔG below 1 kcal/mol, and compared the results with ClinSig and AF matrices. If no significant contradictions were evidenced, we proceed to reclassify the motifs. Motifs in S1f which had variants assigned as Pathogenic or Benign (from either ClinVar or AF), but couldn’t be analyzed in the previous step due to a lack of information, were now considered. Again, if a given motif SAS is classified as benign but in a non tolerant position or pathogenic in a tolerant position, the motif is assigned to N2. While if motif variants classification agree with the ΔΔG matrix they are assigned to P2. Remaining motifs, which could not be classified correspond to S2. As in the previous classification round, the incorporation of new motifs to the positive control group allowed to iterate the process due to the incorporation of new values to the ClinSig and AF matrices, leading to N2f, S2f and P2f. In the next -final-step, we added to the analysis the flexible position of the motif and refined those cases were the substitution has a moderate free energy change, defined as 1<ΔΔG<2 kcal/mol. As for the other positions, we filtered the initially predicted motifs with this new information, assigned them in either positive (P3) or discarded (N3) sets, and added this information to build the definitive SAS tolerance matrix. With this new matrix we can determine which SAS are tolerated or can lead to the motif disruption. Moreover, this matrix allows to determine the flexibility of each position and consequently, better characterize the actual motif consensus sequence.

Having described the general process we turn our attention to a particular case: the phospho-independent tyrosine based motif.

### Phospho-independent Tyrosine-based motif

Tyrosine-based motifs (NPxY) play important roles mediating and regulating the endocytosis of several proteins (Bonifacino and Traub, 2003). The NPxY motifs are recognized by phosphotyrosine-binding (PTB) domains involved in signaling networks as critical scaffold/adaptor proteins (Uhlik et al., 2005). Although, PTB domain basically binds to phosphotyrosine (pY) containing peptides, a subfamily of PTB domains--the Dab-like domains--recognize peptides having non phosphorylated tyrosines or even phenylalanines in the context of the motif (Uhlik et al., 2005). Therefore, our phospho-independent Tyrosine-based motif (PiTb) is defined by the following consensus sequence NPx[Y/F].

We searched in the human proteome the presence of NPx[Y/F] motifs and found 1823 potential motifs (S0) in 1584 proteins. From these, only 15 proteins (P0) were documented and experimentally tested to have functional unphosphorylated tyrosine-based motifs that interact with PTB-domains (Zhang, 1997; Eigenthaler et al., 1997; Trommsdorff et al., 1998; Dho et al., 1998; Homayouni et al., 1999; Su et al., 2002; Calderwood et al., 2003; Marzolo et al., 2003; Stolt et al., 2003; Zhang et al., 2007; Huang et al., 2009; Park et al., 2010; Barbagallo et al., 2011; Fuchigami et al., 2013). Among them, the NPx[Y/F] motifs of Amyloid-beta precursor protein (APP, UniProtID: P05067), Low-density lipoprotein receptor-related protein 8 (LRP8, UniProtID: Q14114), Krev interaction trapped protein 1 (KRIT1, UniProtID: O00522) have been crystallized with their respective PTB domains present in the X11, GULP1, Dab1, ICAP1 and CCM2 proteins (Stolt et al., 2003; Zhang et al., 2007; Liu and Boggon, 2013; Fisher et al., 2015; Chau et al., 2019). According to the corresponding crystal structures, the motif is always located after a beta-extended structure forming a short helix or a loop where the residues Asn and Tyr/Phe of the motif are located in pockets of the PTB domain (Fig 2A). All P0 proteins are localized in the plasma membrane or in the cytosol, the residues that constitute the motif are highly conserved, the motif is preferentially located in the second half of the protein, and in poorly structured or unstructured regions (Fig 2B-D).

**Figure 2.**
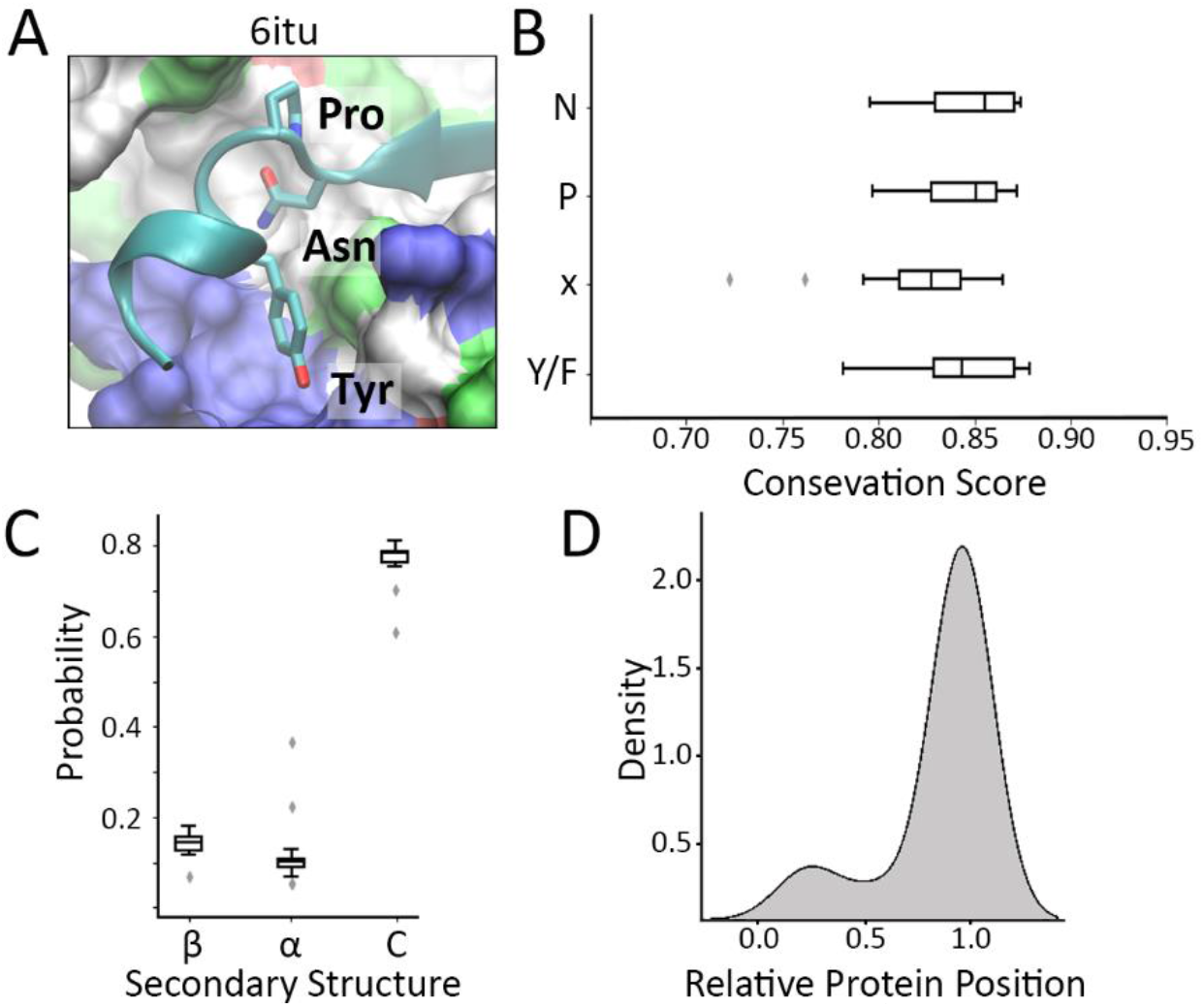
Experimentally tested phospho-independent tyrosine-based motifs. **(A)**. Representative crystal structure of the tyrosine-based motif and Dab1-PTB domain complex. PTB-domain is colored by residue type. **(B)**. Box-plot showing the Jensenn-Shannon Conservation Score of each motif residue. **(C)**. Motif secondary structure prediction. **β**: Beta-sheet. **α**: alpha-helix. **C**: Coil. **(D)**. Histogram representing the distribution of the relative motif position with respect to the whole protein.

As previously described, using the P0 group we constructed the ClinSig and AF Matrices (Fig 3A and B), as well as the ΔΔG matrix using the available crystal structures (Fig 3C).

**Figure 3.**
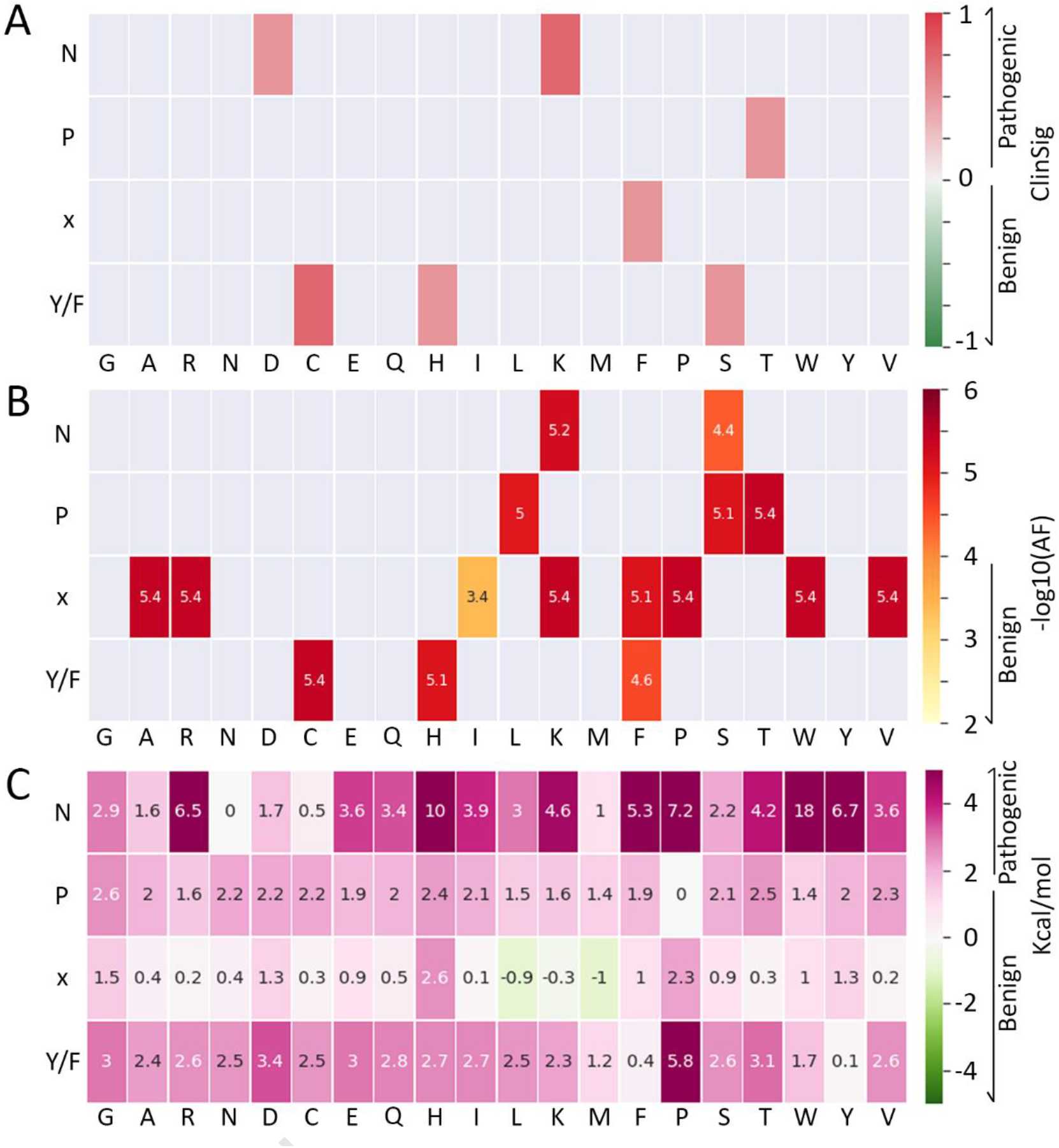
Tyrosine-based motif P0 SAS Matrices. **(A)** ClinSigMatrix. Pathogenic variants are presented in red palette and Benign in green palette. **(B)** AF Matrix. SAS with −log10(AF) < 4 (Bening) are colored in yellow. **(C)** ΔΔG matrix (in kcal/mol). Values > 2 kcal/mol were considered as pathogenic, while values below 1 kcal/mol were considered as probably tolerated for the interaction.

The first classification round (filtering matrix based on ClinSIg and AF data, Fig 4A), allows to discard 22 motifs (N1). As an example, we discarded the transmembrane protein 43 (Q9BTV4) due to the presence of a Benign Tyr233Cys variant (rs35924492) in the predicted motif, which has been reported as Pathogenic (Davis et al., 1986). We also discarded the protein Pro-neuregulin-3 (P56975) since the variant Asn576Lys (rs17101193) has a −log10(AF) < 4, and thus was considered Benign; however, positive charged residues at this position in the motif have been reported as pathogenic (Fig 3) (Etxebarria et al., 2015). This classification round also results in 3 putative proteins that harbor a real tyrosine-based motif: Ephrin-B1 (P98172), SCN1A (P35498) and HUWE1 (Q7Z6Z7). However, analysis of the data in the UniProt database, allows discarding all three, thus moving them also to N1f,because P98172 and P35498 have the motif in an extracellular domain, and Q7Z6Z7 resides in the nucleus. Neither of these features is compatible with the nature of tyrosine-based motif (Bonifacino and Traub, 2003).

**Figure 4.**
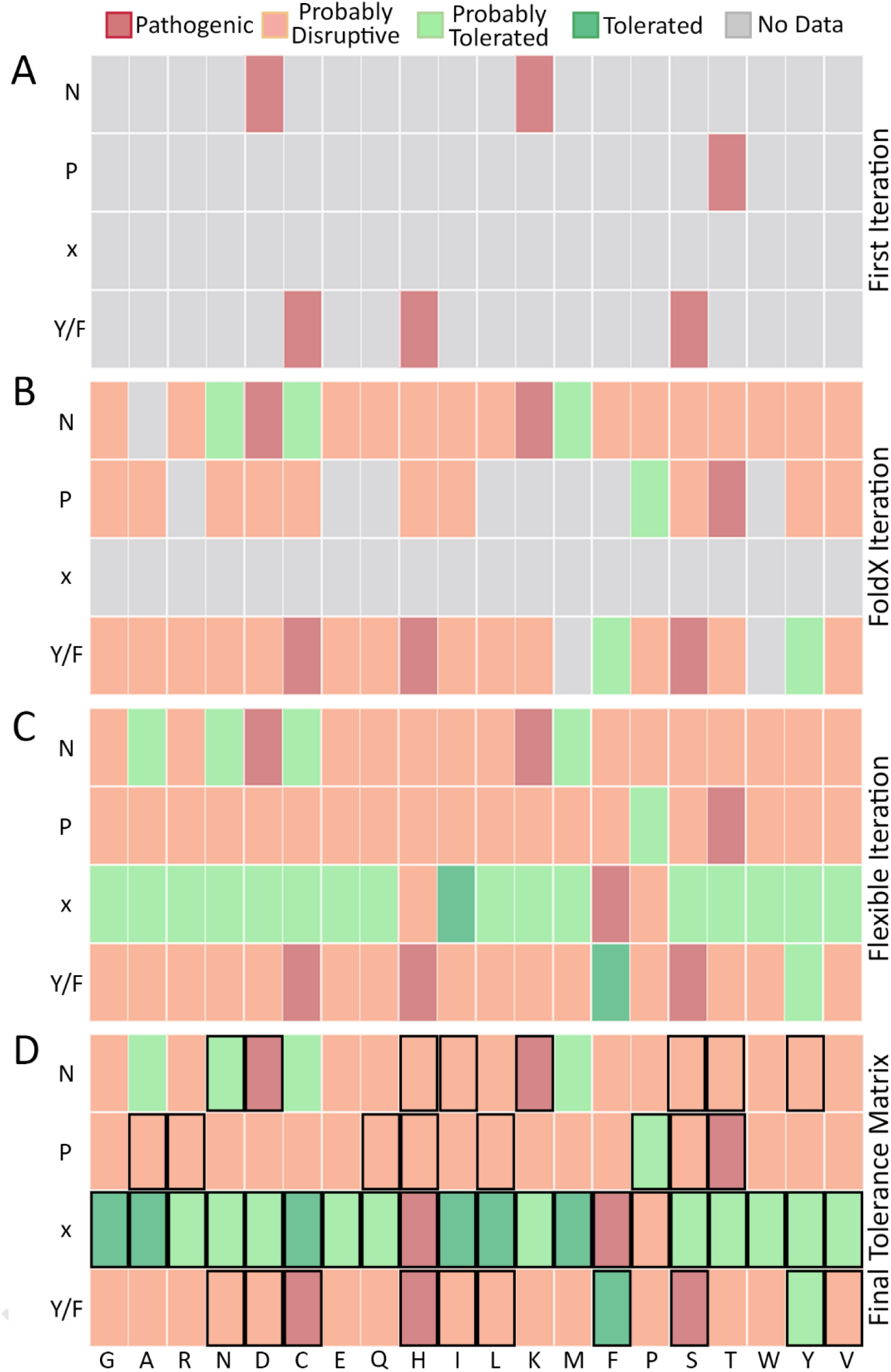
Tyrosine-Based motif SAS matrices. **(A)** initial SAS tolerance matrix based on ClinSig and AF **(B)** 2nd Tolerance matrix with incorporation of ΔΔG matrix and motif classification round. **(C)** 3rd Tolerance matrix including flexible position and refinement of FoldX SAS **(D)** Final SAS tolerance matrix. In all matrices grey rectangles represent those substitutions that were not considered in each filtering step. Black rectangles indicate those SAS that can be the output of Single Nucleotide Variants (SNVs). Dark colors indicate those variants from ClinVar and GnomAD databases.

Classification based on the structural-derived matrix (Fig 4B), allows to discard 61 additional motifs and moving them to N2, since for example Olfactory receptor 8J3 (Q8NGG0) presents a Benign substitution Asn57Thr (rs1947924) but this change has an average ΔΔG of 4.2 kcal/mol and so was considered disruptive. Similar reasons apply for the other cases. We identified 3 proteins that may have a functional motif: DNAH12 (3rd motif) (Q6ZR08), TMCT4 (Q5T4D3) and GALAC1 (P07902). We discarded TMCT4 since it is an ER-resident protein with the motif located in a luminal region. GALAC1 is a Galactose-1-phosphate uridylyltransferase localized in the cytosol. All the features of the predicted motif in this protein are in accordance to the positive control, and thus, we predict with high confidence that this protein may have a functional tyrosine-based motif at the protein position 86-89. This region of the protein is highly unstructured as revealed by the x-ray-solved 3D structure (McCorvie et al., 2016). The mutations Y89H (rs111033666) and Y89D (rs111033666), that affect the predicted tyrosine-based motif have been linked to Classical Galactosemia (Singh et al., 2012). Clearly there are strong arguments as this being a functional motif. However, it is also important to note that functional characterization of the Y89H mutant revealed that only 15% of the protein activity is conserved, suggesting the substitution may affect protein stability (Singh et al., 2012). We analyzed the impact of both reported mutations at position Y89 to the protein stability using FoldX, showing that both present values of > 6kcal/mol, thus also suggesting that protein structure is affected. Therefore, although the data clearly shows that mutations in this position are not tolerated, we cannot ascertain whether it is due to the presence of a functional motif, due to production of an unstable protein, or both. Finally, the protein DNAH12 has a Benign Tyr2740Phe variant (rs17057989) affecting the third predicted tyrosine-based motif. This region of the protein is conserved and unstructured, in agreement with the characteristics of a functional motif. Thereafter, we added it to the positive control group.

Afterwards, we integrated the information of the flexible position and of some substitutions that were not considered previously (refinement) in the next SAS tolerance matrix (Fig 4C). With this information, we discarded 27 additional motifs and we identified 44 new candidate motifs. We discarded 8 due to low conservation scores (Q7Z4L5 (2nd motif), P01160, Q96PV0, Q9GZU1, Q6ZR08 (2nd motif), Q5HYI8, Q32MH5, P41161), 6 because they present secondary structures within the motif (Q96RY7, P20807, Q8NH48, Q9UHA7, Q6QHF9, Q86YT5), 5 due to their presence in the N-terminal section of the protein (Q8WV16, Q9H7Z3, Q6PP77, Q96PV0, P20807) and 14 taking into account the UniProt data (P16066, P16989, P09017, Q86×52 (2nd motif), P56470, Q6NUN7, Q9Y4A5 (2nd motif), Q5T764 (2nd motif), Q99594, Q8N2K0, P55210, Q02763, Q9UBX5, Q8TAB3).

We finally retained 10 proteins that may have real tyrosine-based motifs (Q8IX94, P0CG41, Q13635, P26010 (2nd motif), Q9BSA4, Q5T7P8 (1st motif), Q9P2P5, Q9HAI6, Q14183, Q8NHH9) and incorporated them in P3f. All of these show no contradiction and one variant whose characteristics agrees with the tolerance matrix. The only protein that has more than one variant congruent with the SAS tolerance matrix, and thus higher confidence is Q9BSA4. Other minor points of note are protein E2 ubiquitin ligase HECW2 (Q9P2P5) which has a pathogenic substitution Tyr1277His (rs1575255333) (flexible position of the motif) reported in ClinVar, but there is no functional evidence that this mutation produces the disease, and the integrin beta-7 (P26010) which has a phospho-dependent tyrosine based motif. In the last step, we combined all the information to build the final SAS tolerance matrix (Fig 4D). Finally, we classified 158 motifs: 147 motifs are non-functional and 11 have high confidence of being biologically relevant (S1 Table). Only 2 motifs that present SAS with either ClinSig or - log10(AF)<4 could not be classified due to the lack of UniProt information.

The final SAS tolerance matrix serves three purposes. First, as shown by the previous filters, it allows better prediction as to whether a given motif, found using regular expression, is (or not) functional. Second, it delivers a more detailed description of the motif. In the present example, the initial description of the Tyrosine-based motif is that it displays the form NPx[Y/F]. This is confirmed for all three well defined sites, where only specific changes could be additionally tolerated, such as Ala, Cys or Met in the first position, and perhaps Trp or Met in the 4th position. Most important, our matrix shows that not all 20 residues are tolerated in the x position, since His, Pro and Phe seem not to be tolerated in the peptide-complex interaction (Fig 4D), information that allowed us to discard 194 additional motifs. Finally, the SAS matrix is expected to aid in the classification of new variants, since those corresponding to non tolerated SAS would argue in favor of their being pathogenic, while those in tolerated SAS positions give support to their benign nature.

We applied the same procedure to others SLiMs derived from the ELM database. In the following sections, we briefly describe the final positive and negative sets and SAS tolerance matrix for each of them. The ClinSig, AF and ΔΔG matrices, together with UniprotIDs of potentially new proteins harboring the motifs as well as those discarded are available in SI. The first presented case which corresponds to Type 1 PDZ-binding motif, was also used to perform a preliminary validation of the proposed methodology. We selected this motif since it has a relatively large positive control group with 71 known and validated motifs and so, a good amount of motif occurring variants.

### Preliminary validation of the SLiM identification pipeline using Type 1 PDZ-binding motif

PDZ-binding motifs are located at the carboxy-terminus, and are recognized by abundant protein interaction modules named as PDZ domains (Harris and Lim, 2001; Sheng and Sala, 2001). The different classes or types of PDZ-binding motifs differ in the nature of residues that constitute the motif and therefore, the type of interaction that takes place with the PDZ domain. The type 1 PDZ-binding motif ([S/T]x[V/I/L]-COOH) is one of the most widely studied motifs in the human proteome. The interaction between type 1 PDZ-binding motifs and PDZ domains determine the stability and subcellular localization of the target protein (Ivarsson, 2012). PDZ-binding motif-containing proteins are mainly located at the plasma membrane, but also in the nucleus, membranous organelles and cytosol. The human proteome shows the presence of 1022 motifs from which 71 were documented (and experimentally validated) of having a functional type 1 PDZ-binding motif. Over 15 structures of peptides containing a type 1 PDZ-binding motif in complex with PDZ domains have been reported in the PDB. The backbone of the peptide containing the PDZ-binding motif binds to the PDZ domain via β augmentation (it forms a β-strand with a neighbouring β-sheet in the domain), while the side chains of the ligand residues interact with a neighbouring α-helix in the PDZ domain (Ivarsson, 2012). For this interaction to take place, the PDZ-binding motif has to be initially unstructured, and then adopts the β-strand configuration in the complex (S2 Fig). The classic type 1 PDZ-binding motifs are carboxy-terminus located motifs, however there are some reports that revealed the presence of internal non-canonical PDZ-binding motifs (Ivarsson, 2012). Here, we only considered the canonical carboxy-terminus PDZ-binding motifs.

To have a preliminary assessment of our proposed method capacity of detecting with high confidence new (i.e previously unknown or non validated) motifs, we performed the following analysis. From the 71 known PDZ binding motifs, we found 16 which have at least 1 variant classified as pathogenic or benign due to its ClinSig or AF. We therefore divided these 16 motifs randomly in two parts and assigned one half to P0 and the other to S0. The P0 was also complemented with the general information of the remaining positive motifs. Subsequently, we applied the whole pipeline and looked at how many of the known motifs assigned to S0 were retrieved in P3f.

Using the first P0 set, we were able to correctly predict all eight type 1 PDZ-binding motifs that were experimentally validated. The true positive motifs were all predicted during the FoldX or Flexible iteration step. Furthermore, with this (set 1) tolerance matrix we discarded 25 motifs and predicted with high confidence 23 additional motifs (16 with low confidence) of being a functional PDZ-binding motif. Using the second P0 set we were able to correctly predict 7 out of 8 known type 1 PDZ-binding motifs. Again, for set 2, the true positive motifs were predicted during the FoldX or Flexible iteration step. Also, we discarded 32 motifs and predicted with high confidence 20 additional motifs (13 with low confidence).

The false negative corresponds to PDZ-binding motif (P43250), and is discarded due to the reported Thr574Pro benign substitution (rs763634515), which our tolerance matrix assigns as non tolerable. Interestingly, this SAS also produces the only observed contradiction between the GnomAD and FoldX matrix (using the whole true positives) (S3 Fig). The FoldX matrix shows a ΔΔG > 6 kcal/mol for this amino acid change, but displays −log10(AF) value of 2.4, which we assign as benign. Interestingly, other proteins of the whole positive control group present substitutions of Ser/Thr by Pro with low AF (-log(AF)>4), such is the case of the Thr442Pro variant (rs1300757305) of the Corticotropin-releasing factor receptor 1 (P34998), or the substitution Ser4653Pro (rs748754632) of the Low-density lipoprotein receptor-related protein 2 (P98164). In this scenario, we believe it is more likely that the variant is actually pathogenic, despite its moderate frequency in one protein, and the tolerance matrix should show Thr to Pro as a non tolerant SAS in the motif first position. In any case, and despite the above mentioned detailed analysis, our preliminary cross validation showed that we were able to retrieve 15/16 true motifs, which we consider a very good performance.

Taking these considerations into account and using the whole positive dataset, we discarded 22 motifs and predicted with high confidence 25 new motifs (S1 Table). Furthermore, 18 motifs show no SAS contradiction but lack UniProt information for assessing its biological relevance. The final tolerance matrix (Fig 5A) revealed that only Gly and Pro are not tolerated at the initial position while all the aminoacids seem to be well tolerated at the flexible position. The last position, that constitutes the core of the PDZ-binding motif - PDZ-domain interaction, can tolerate a Met in addition to the Leu, Ile and Val, but not Ala, Cys or Phe as suggested in the ELM resource.

**Figure 5.**
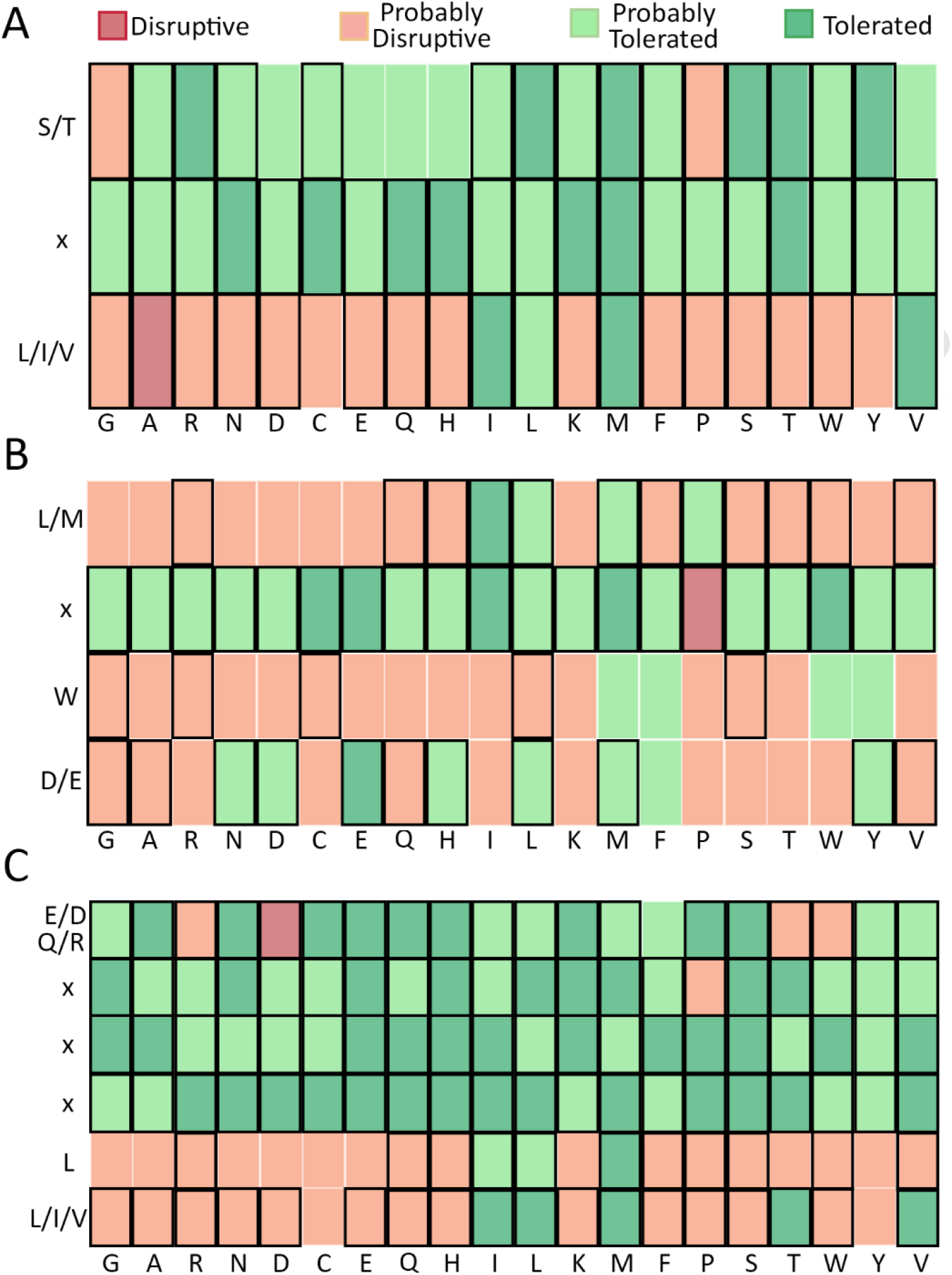
Final SAS matrices. **(A)** Type 1 PDZ-binding motif final SAS tolerance matrix. **(B)** Tryptophan-acidic motif final SAS tolerance matrix. **(C)** Acidic di-Leucine motif final SAS tolerance matrix. Black rectangles indicate those SAS that can be the output of Single Nucleotide Variants (SNVs). Dark colors indicate those variants from ClinVar and GnomAD databases.

### Tryptophan-acidic motif

Tryptophan-acidic motifs ([L/M]xW[D/E]) are recognized by the light chain (KLC) of the molecular motor complex Kinesin-1. KLC interacts with tryptophan-acidic motifs via tetratricopeptide repeat (TPR) domains, regulating the microtubule-based cargo trafficking and hence, its final subcellular localization (Dodding et al., 2011). Some tryptophan-acidic motif-containing proteins may have the motif in a structured conformation that lately change and adopt an unstructured conformation in order to enable the motif-TPR domain interaction (Wilson and Holzbaur, 2015). The human proteome analysis retrieved 1894 potential motifs, of which only 8 are documented (and experimentally validated) of having a functional tryptophan-acidic motif. Only 2 crystals of peptides containing a tryptophan-acidic motif in complex with TPR domains have been reported. The structure, shows that the tryptophan-acidic motif-containing peptide has a completely unstructured conformation, where the Leu and Trp residues present hydrophobic interactions with pockets of the TPR domain, while the acidic residue makes polar interactions with the amino acids located on the surface of the TPR domain (Pernigo et al., 2013). These proteins that constitute the positive control are localized at the plasma membrane, membranous organelles, nuclear membrane or in the cytosol, the residues that constitute the motif are highly conserved, the motif is preferentially located in the second half of the protein and in poorly structured or unstructured regions (S4 Fig).

After the iterative classification process, we were able to discard 102 motifs and identify 10 new high confidence potential candidates, and one additional potential motif which shows no SAS contradiction, but lacks information in UniProt (S1 Table). Analysis of the SAS tolerance matrix (Figs 5B and S5) shows that the first position, in addition to L/M allows the presence of Ile and perhaps even Pro. The third position, W which also seems to accept the other aromatic residues (Y/F) and even Met. The fourth acid (D/E) position, possibly tolerates other polar residues such as Asn, His or Tyr and some neutral residues such as Leu, Met or Phe. Finally, the flexible position seems to tolerate well all residues except Pro. It should be noted however, that the low amount of ClinSig variants makes these predictions to have moderate confidence.

### Acidic di-Leucine motif

Leucine based motifs have been widely studied in intracellular trafficking, particularly in polarized sorting and endocytosis (Bonifacino and Traub, 2003). The nature of the residues in the vicinity of the “di-Leucines’’ determine the type of interaction that will take place. Acidic di-Leucine motifs are known interaction partners of GGAs proteins (Golgi-localizing, gamma-adaptin ear homology domain, ARF-binding proteins) and heterotetrameric Adaptor Proteins (APs) (Bonifacino and Traub, 2003). Those motifs defined by the consensus sequence [D/E]xxL[L/I/V] are recognized by the VHS domain (Vps-27, Hrs and STAM) of GGAs, while those defined by [D/E/Q/R]xxxL[L/I/V] motifs interact with a pocket conformed by sigma and alpha (AP1) or sigma and gamma (AP2) clathrin adaptor complex’s subunits. Due to the regular expressions overlap, and therefore, some of these motifs interact with both GGAs and Aps (Doray et al., 2008).

For our study we centered our attention on those acidic di-Leucine motifs that interact with APs. Despite its extension in the human proteome (47821 potential motifs in 15376 proteins), only 20 motifs have been experimentally validated, and only 4 crystal structures of the corresponding complexes have been documented. The crystal structures revealed that the di-Leucine comprehends the core of the interaction between the motif and the AP. Particularly, these hydrophobic residues interact with a pocket of the sigma subunit. The polar residues located downstream interact with the alpha (AP1) or gamma (AP2) subunits, however this interaction seems no to be essential for motif recognition. The acidic di-leucine motifs are highly conserved, displayed in unstructured regions and preferentially located near the amino or carboxy-terminus (S6 Fig). Proteins containing these motifs are transmembrane proteins located at the plasma membrane or membranous organelles.

After the iterative classification process, we were able to discard 6285 motifs and identify 171 new high confidence potential candidates, 940 additional candidates lack information in UniProt and therefore should be considered as awaiting further information (S1 Table). Surprisingly, the SAS tolerance matrix (Figs 5C and S7) shows that the first position seems to not tolerate the presence of Arg, Asp, Thr and Trp, despite having Arg and Asp as part of the consensus sequence. It should be noted however, that these results may be biased due to the crystal structures used and the lack of ClinSig variants. The more restrictive positions are the last two that comprehend the core of the interaction between the motif and the adaptor protein. The fifth position can only tolerate Ile and Met, while the last position can tolerate Met and Thr in addition to Val, Ile and Leu. The flexible positions can tolerate any amoniacid, except for Pro at the second position of the motif. This information allowed us to discard 2225 additional motifs.

## Discussion

SLiMs are central elements in protein physiology and cellular biology, since their presence controls regulation and subcellular localization of the motif-containing protein. SLiMs are usually located in poorly structured and solvent-accessible protein regions, which enables motif recognition by the corresponding interaction partners. This protein-protein interaction determines the proper functioning of the cellular protein network and consequently, the cellular homeostasis. Knowing the SLiMs that mediate cellular pathways can have significant outcomes, such as blocking the internalization and replication of SARS-CoV-2, as the Angiotensin-converting enzyme 2 (ACE2) and some integrins (molecules that also act as receptors of the virus) have been shown to contain many SLiMs mainly involved in endocytosis (tyrosine-based motifs) and autophagy (LC3-interacting region motif) (Mészáros et al., 2021). On the other hand, disruption of the cellular protein network leads to improper cellular functioning and therefore, to a disease development. SLiMs identification softwares usually query protein sequences for the presence of motif consensus sequence or regular expression pattern. Many use additional logical based filters, conservation scores and, more recently, Hidden Markov models or machine learning techniques, in order to improve their predictability.

However, only a fraction of the predicted motifs are biologically relevant. Therefore, precise functional SLiMs determination remains a challenge. In the present work, we show that motif identification can be significanlty improved by the inclusion of information derived from intraspecies variation, i.e human genome variant frequency and Clinical Significance, combined with structure based interaction energy calculations between the SLiM and its receptor.

For each SLiM we iteratively built a SAS tolerance matrix, which reflects in each motif position whether a given substitution is tolerated or disruptive. We started by looking at ClinSig information, and assumed that if a SAS in a previously established motif is shown to be classified as clinically benign it should be tolerated and if classified as pathogenic it is not. As shown in Table 1, the number of true experimentally proved motifs with respect to those that can be found using regular expressions is very small. Close, or even smaller than 1%. Moreover, the number of motifs with reported variants in ClinVar is also very small, thus our initial ClinSig matrices were almost empty. Inclusion of variant frequency information, improved our tolerance matrices completeness, but still not enough. The key step, was to use each motif-domain crystal structures to quantify each SAS ΔΔG, and finally to add each new high confident non contradictory motif information (ClinSig and AF) to the matrix.

At this point it is important to remark that obtention of a high confidence SAS tolerance matrix, requires careful observation of positive feedback (i.e coincidence) of ClinSig, AF and FoldX matrices, and resolution of those small number of positions which show conflicting interpretations. For example, substitution tryptophan-acidic motif Leu at position 2, by Ile shows a moderately high ΔΔG value of 2.8 kcal/mol, which could indicate this change is not tolerated. However, analysis of the AF of the corresponding variants undoubtedly classifies it as benign, which together with analysis of the available information for those motifs displaying this SAS confirms that the change is tolerated. Another, opposite example, is presented by the PDZ-binding motif, and the case that leads to a false negative prediction when performing the previously shown cross-validation. Here, the FoldX matrix shows that Thr to Pro substitution is not tolerated. However, one protein with a functional motif shows this variant with relative high AF, which could be classified as benign. Interestingly, in other proteins the same variant displays significant lower AF, consistent with a non tolerable change. In this case, our conclusion is that the SAS should be considered as non tolerable, and that the variant is possibly pathogenic, even though it has a moderate AF. These examples underscore the relevance of our strategy which combines the information for each SAS from three different and independent sources (ClinSig, AF and Structure), and also suggests as key issues the careful analysis of the final tolerance matrix to properly solve inconsistencies.

In addition to their predictive power (see below), obtention of the final SAS tolerance matrix also results in a deeper understanding of the motif and its possible sequence variations. For example, as seen for the phospho-independent tyrosine-based motif, the presence of His, Pro or Phe at the flexible position seems to alter the motif recognition by its corresponding domain (Fig 4D) and thus are avoided. Moreover, the presence of Pro is also not tolerated in the first flexible position of the acidic di-Leucine motif and the flexible position of the tryptophan-acidic motif (Figs 5B and C) underscoring the relevance of considering the impact of each SAS at the flexible positions, and showing that although flexible not all residues are tolerated. Another interesting topic is the expansion of amino acids that can take place at motif defining position according to the Consensus Sequence or Regular Expression which, in general, constitutes the interaction. Such is the case of the presence of Met at the last position of type 1 PDZ-binding motif and acidic di-leucine motif (Figs 5A and B).

These results suggest that the consensus sequence of each motif should be re-considered with corresponding impact in the universe of proteins that may have a functional motif. For example, it is interesting to mention, that for some motifs, we found a discrepancy between the final tolerance matrix and the regular expression (or consensus sequence). Particularly the case of the acidic di-leucine motif where the first position has been proposed to tolerate Glu, Asp, Gln and Arg, but our final tolerance matrix shows that neither Arg nor Asp can be tolerated at this position (Fig 5C), while other non-charged amino acids seems to be tolerated. In this case, we believe that our results could be biased by the available crystal structures used for the analysis where the motifs only contain Glu at the first position. In any case, the point we want to make is that building the tolerance matrix and analysing it in the context of the motif original regular expression, can help to refine the motif definition and thus its subsequent searches in the protein universe.

Concerning our strategy, predictive capacity numbers are promising. Our preliminary validation with Type-1 PDZ-binding motif showed that we could correctly retrieve 15 out of 16 true positive motifs. Generally speaking, in all cases we could identify a significant number of novel motifs with high confidence, the numbers represent 2 to 10 times increase in the original true positive set depending on the motif analyzed. Most importantly we were able to discard a significant number of motifs, also with high confidence. We are able to classify with high confidence all motifs included in the S0 set, i.e those where variants have been found (either in ClinVar or in GenomAD). The S0 set currently represents 10-20% of the initial potential motif set. However, as more and more human genomes and exomes are sequenced and uploaded to public databases this number increases faster and faster. Therefore, our method predictive capacity, both in terms of precision and number of motifs analyzed, also grows continually.

Another point of notice, concerning future perspectives, concerns the current applicability scope of the presented strategy. To apply our method, we need to: i) have an initial set of known true positive (exp. validated) motifs (usually about 10) with ClinSig or AF information, ii) have at least one crystal structure of the motif-receptor complex to build the FoldX matrix and iii) have a moderately large set (over 100) of proteins harboring a potential motif with reported variants (i.e S0). To have a preliminary assessment of the potential scope of the presented method, we surveyed the whole ELM database. Our results (Table 2), show that in addition to the presently analyzed motifs, there are 27 additional motif classes (out of 296) that have at least 10 true human motifs, and at least one crystal structure of the motif-receptor complex. By analyzing the number of motif-occurring variants we discarded 2 motifs due to the lack of ClinSig variants in the motifs that constitute the positive control. Finally, 25 motifs are clear candidates for near future application of the herein presented strategy. Moreover, it is important to note that in our experience, manual literature search usually increases the number of true positive motifs with respect to those reported in ELM and helps to refine the motif Consensus Sequence, thus likely increasing method accuracy.

**Table 2.**
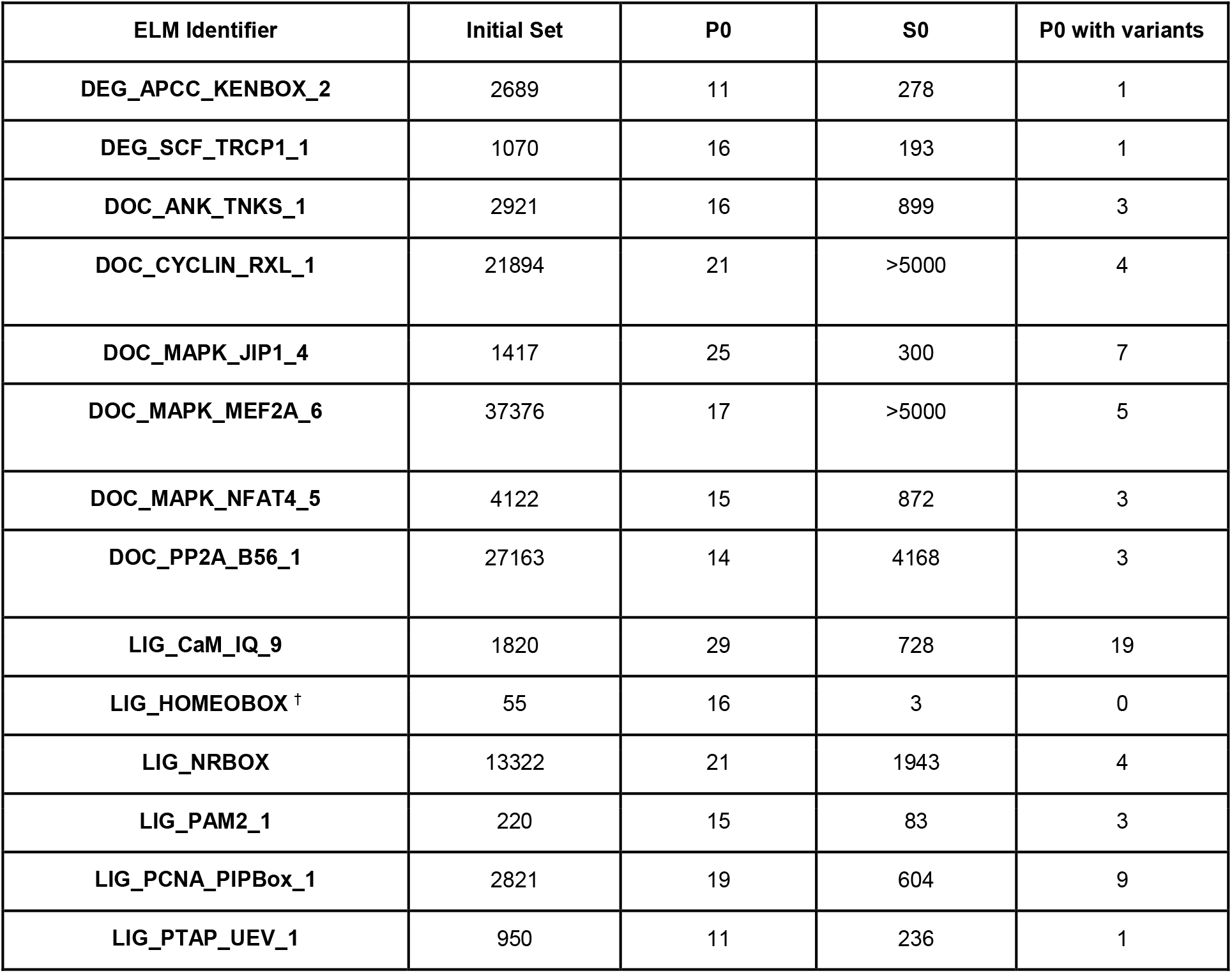
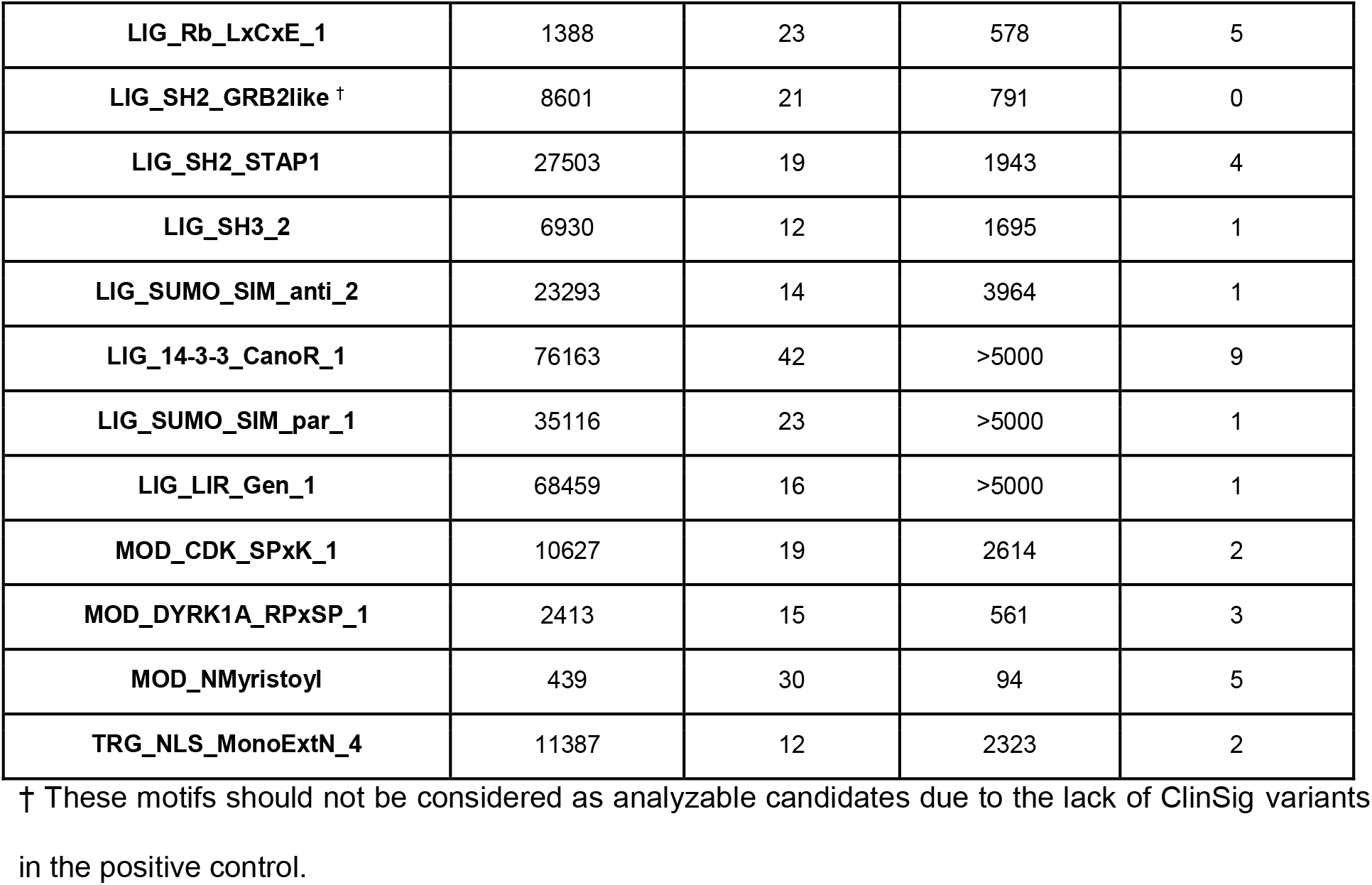
SLiM candidates with initial data information.

Another potential source of seeds for our method is the Prosite database, which also presents regular expression of motifs with the corresponding MSAs and identified human proteins. Although the present work was focused on motifs related to protein trafficking we expect to be able to analyze its applicability to other types of motifs, such as those involved in post translational modifications where the receptor is an enzyme and the motif key part of its substrate.

Finally, the proper constitution of motifs substitution matrices is also important for the prediction of new Clinical Significance of variants occurring within the motif. According to our analysis, over 70% of the SAS that can occur due to SNVs in a motif “rigid” position are predicted to be disrupting and consequently, may lead to a pathogenic phenotype. If we add the percentage of non tolerated SAS at the flexible position this percentage is still around ∼40-50%. In other words, the reverse side of the coin, of using variant information to predict true functioning motifs with high confidence, is the use of our knowledge of high confidence identified motifs to better classify novel variants leading to SAS inside them. For a real working example, consider that currently, ACMG guidelines assign PM1 (Pathogenic Moderate 1) to those variants found in a well studied functional domain, without benign variants. If taken strictly, most SAS inside SLiMs will not fulfill this criteria, since either the SLiM is not confirmed, or even if its, as shown here, usually benign variants can be present, specially in the flexible position. However, using our data, we could refine PM1 assignment when: i) the motif is predicted as a high confidence motif and ii) the variant corresponds to a SAS in a non tolerated position. Although other ACMG criteria are not as straightforward applicable to SLiMs, knowledge of the presence or absence of a particular SLiM with high confidence and analysis of a novel SAS in light of the final SAS tolerance matrix, definitely adds relevant information for variant interpretation and classification, thus potentially leading to better diagnostics.

Last but not least, we want to mention two current drawbacks of our proposed strategy. The first, is that given that there is no ClinSig or AF information for non human species, our method is restricted to human SLiMs. However, the rationale of the presented method could be adapted to other species, provided experimental information akin to ClinSig or AF can be produced, for example, using massive mutation selection experiments in microorganism (bacteria, yeast) and/or using species (or higher taxa) population genetic data.

Another drawback concerns the potential impact of SAS in regions adjacent to the SLiM that significantly change its receptor recognition Accordingly, we recently showed that the variant G561E in the sodium/iodide symporter, preceding a tryptophan-acidic sorting motif in the carboxy-terminus of the transporter, shifts the equilibrium of the unstructured tryptophan-acidic motif towards a more structured conformation unrecognizable by KLC2, thus leading to congenital hypothyroidism (Martín et al., 2021a). Although our strategy currently is strictly limited to SLiMs regular expression defined sequence, the same procedure can be extended to adjacent residues, provided there is variant and structural information available.

The rise of massive sequencing techniques and their implementation in the clinic, the increasing experimental validation of predicted SLiMs and new ligand-domain crystal structures will allow to improve the construction of substitution matrices that reflect the real biological effect of each amino acid change, leading to a better understanding of the effect of amino acid substitutions within a given motif and consequently reaching to more precise and accurate predictions of biological relevant SLiMs.

## Methods

### Regular Expression and positive controls for SLiMs

Regular Expressions are written according to ELM database nomenclature (Kumar et al., 2019). Previously characterized SLiMs used as positive controls are referred using the corresponding UniProtID (The UniProt Consortium, 2021).

#### - Phospho-independent Tyrosine-based motif

Regular expression: NP.[YF]

Consensus sequence: NPx[Y/F]

ELM Identifier: LIG_PTB_Apo_2

Positive controls: P01130, P05067, Q14114, P05556 (1st and 2nd motif), Q8TBB1, Q07954, Q9NY15 (2nd motif), P05107, P05106 (2nd motif), P98164 (3rd and 4th motif), P51693, Q06481, O00522 (2nd motif), P98155, Q9HCE7.

#### - Type 1 PDZ-binding motif

Regular expression: [ST].[VIL]$

Consensus sequence: [S/T]x[V/I/L]COOH

ELM Identifier: LIG_PDZ_Class_1

Positive controls: Q8TEU7, P07550, P35222, P46937, Q13268, Q86XX4, Q9H251, P23634, P11274, P48050, P09619, Q495M9, P13569, Q9P0K1, Q01814, Q8TD84, Q96A54, O95477, Q9UQB3, P16473, P32241, O00548, P53778, Q8WXS5, Q01668, Q00722, Q9UBN4, P11166, P36507, Q13936, P34998, Q14524, P47900, P25025, Q8N695, P60484, Q9NQ66, Q9UL62, Q92806, O75096, Q15046, P48960, P25054, Q9Y5I7, P98164, Q9NSA0, O95436, Q5JU85, Q13224, O00192, P30872, P28223, Q04656, P35228, Q9UHC3, Q14916, P43250, P42261, O43424, P23508, Q9Y6M7, P46721, Q96BN8, P08588, P35609, Q9P2W3, O60469, Q9UKN7, O43572, Q99527, Q92911.

#### - Tryptophan-acidic motif

Regular expression: [LM].W[DE]

Consensus sequence: [L/M]xW[D/E]

ELM Identifier: LIG_KLC1_WD_1

Positive controls: Q8IWE5, O94985, Q8WXH0 (2nd motif), Q63HQ0 (2nd motif), Q86WG3, Q8NF91 (2nd motif), Q8N205, Q92911.

#### - Acidic di-Leucine motif

Regular expression: [EDQR]…L[LIV]

Consensus sequence: [E/D/Q/R]xxxL[L/I/V]

ELM Identifier: TRG_LysEnd_APsAcLL_1

Positive controls: P01730 (5th motif), P41143 (1st motif), P56817 (1st motif), P16070 (2nd motif), Q04656 (10th motif), P50895 (5th motif), O15118 (3rd motif), P04233 (1st motif), O75204 (2nd motif), Q9H3U5 (1st motif), P33527 (2nd motif), Q04671 (1st motif) Q9NY64 (1st motif), Q8TD20 (1st motif), Q8NHS3 (1st motif), Q9NRA2 (1st motif), O75379 (1st motif), P14672 (3rd motif), Q14108 (2nd motif), P14679 (3rd motif).

### SLiMs identification in the Human proteome

SLiMs were identified using an in house-developed python script based on basic sequence analysis considering SLiM regular expressions described in the ELM database (Kumar et al., 2019) and/or bibliographical sources. We queried all protein sequences annotated as coded in the *Homo sapiens* genome in the UniProt database (The UniProt Consortium, 2021) containing 20,365 proteins (April 2020), here defined as the whole human proteome. All proteins harboring a sequence segment that matches the regular expression, were retrieved and named “Initial Set”. Additionally, the position of the motif in the protein sequence and the length of the sequence were used to determine the relative position of the motif.

### Identification of naturally occurring variants within each motif

Missense variants corresponding to Single Amino Acid Substitutions (SAS) for each motif were extracted from ClinVar (April 2020 - FullRelease) and GnomAD (May 2020 - Exomes) databases. Only variants classified as either (Likely) Pathogenic or (Likely) Benign were considered from ClinVar database.

### Structure data set

The protein structures were retrieved from the Protein Data Bank (Berman, 2000). For the phospho-independent tyrosine-based motif analysis we used 5 crystals (PDB-ID: 1aqc, 1ntv, 4jif, 4wj7 and 6itu) (Stolt et al., 2003; Zhang et al., 2007; Liu and Boggon, 2013; Fisher et al., 2015; Chau et al., 2019), that have a Dab-like PTB domain in complex with unphosphorylated tyrosine-based motif. For the type 1 PDZ-binding motif analysis we used 15 crystales (PDB-ID: 2qt5, 3gj9, 3k1r, 3lny, 3qgl, 3rl7, 3rl8, 4k75, 4q6h, 4wyu, 5eyz, 5ez0, 6ms1, 6mtv, 6y38) (Long et al., 2008; Yan et al., 2009, 2010; Zhang et al., 2011, 2010; Balana et al., 2011; Amacher et al., 2014; Ren et al., 2015; Maisonneuve et al., 2016; Caria et al., 2019; How et al., 2019; Zhu et al., 2020), that have a PDZ domain in complex with type 1 PDZ-binding motif. For the tryptophan-acidic motif analysis we used 2 crystals (PDB-ID: 3zfw, 6f9i) (Pernigo et al., 2013; Cockburn et al., 2018), that have KLC2 TPR domains in complex with tryptophan-acidic motif. For the acidic di-leucine motif analysis we used 4 crystals (PDB-ID: 2jkt, 4nee, 4p6z, 6uri) (Kelly et al., 2008; Jia et al., 2014; Ren et al., 2014; Kwon et al., 2020), that have subunits sigma and alpha/gamma (AP2 and AP1, respectively) in complex with acidic di-leucine motif.

### Free-energy change calculations

The free-energy change (ΔΔG) of each SAS was calculated using the FoldX software (Delgado et al., 2019) from either one or several structures of the corresponding SLiM receptor complex. The results for each motif are represented in the corresponding free-energy substitution matrix. When multiple structures were available values in the corresponding matrix correspond to the average.

### Protein structure visualization

The structure visualization and figures were made using Visual Molecular Dynamics software and Seaborn Library (Humphrey et al., 1996; Waskom, 2021).

### Secondary structure prediction

Motif secondary structure prediction was performed using sequences of 200 amino acids length, with the motif positioned in the middle. The prediction was performed using the Jpred API (Drozdetskiy et al., 2015), that gives as output the probability of having a beta-extended, alpha-helix or coil conformation in the target sequence.

### Conservation Score calculation

Multiple sequence alignments of target proteins from the UniRef90 cluster were performed with ClustalO software (Madeira et al., 2019) and used as input for calculating motif conservation score. The motif conservation score was calculated using the Jensen-Shannon divergence as previously described (Capra and Singh, 2007).

## Acknowledgements

We acknowledge to Franco Brunello that assisted and discussed the initial project idea.

## Supporting Information

Analyzed motif ClinSig, AF and FoldX matrices, list containing high confidence predicted true motifs and discarded motif (using UniProtIDs) are provided as supplementary information.

**Supplementary Figure 1.**
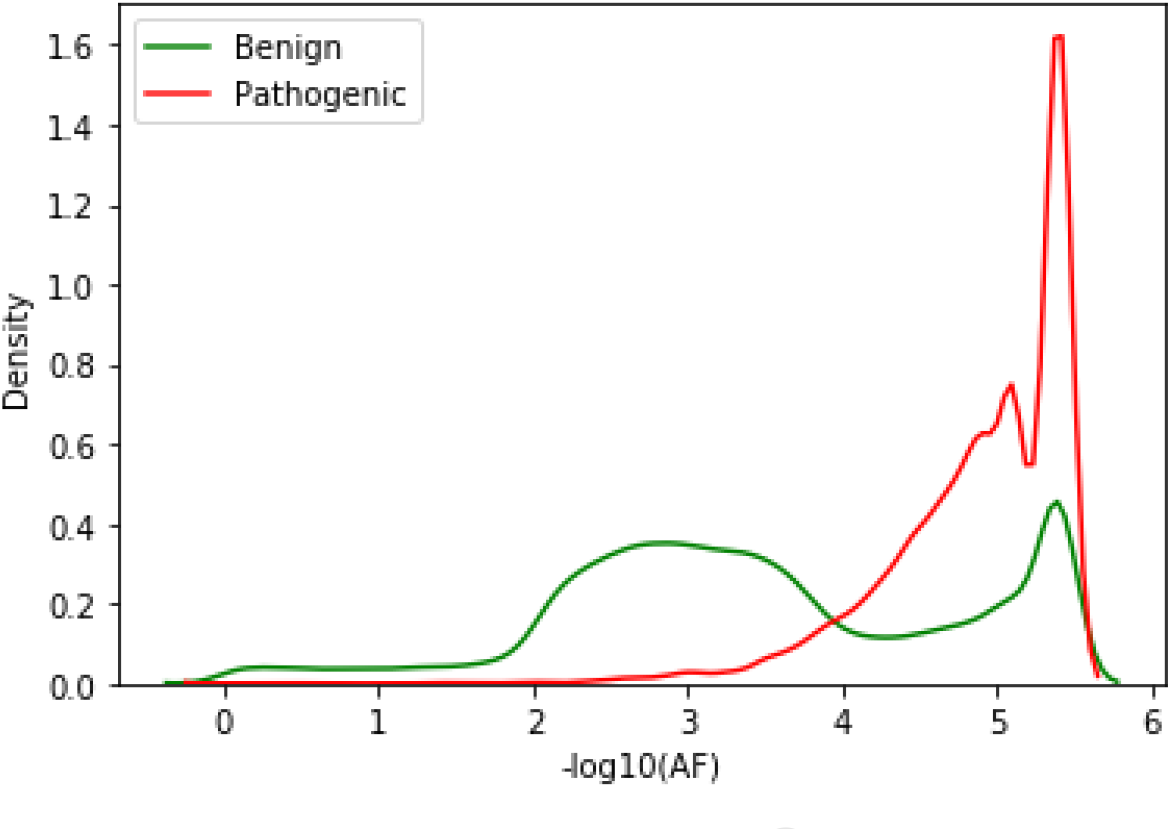
Allelic Frequency distribution of missense variants. Benign variants are represented in green and Pathogenic variants in red.

**Supplementary Figure 2.**
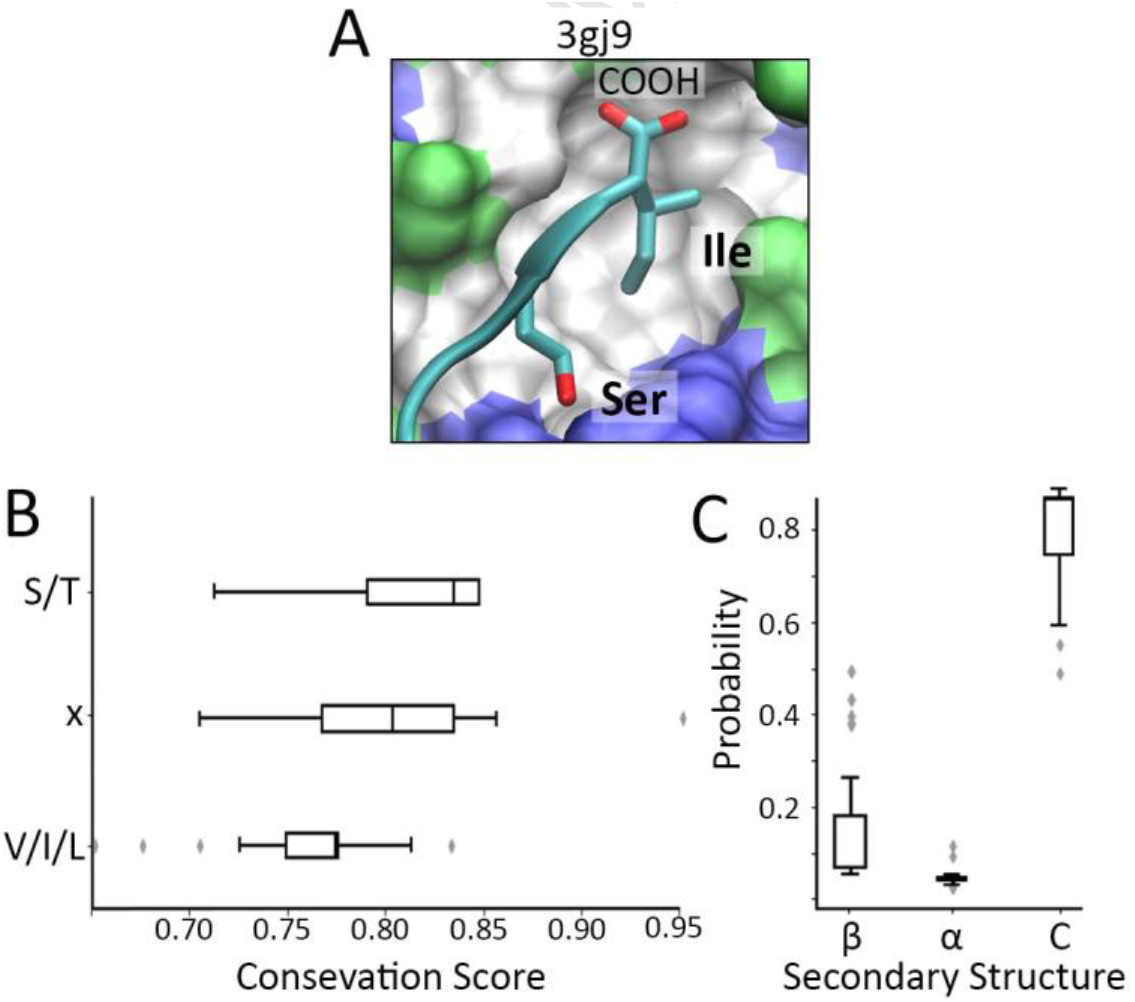
Experimentally tested type 1 PDZ-binding motif. (A). Representative crystal structure of the type 1 PDZ-binding motig and PDZ domain complex. PDZ-domain is colored by residue type. (B). Box-plot showing the Jensenn-Shannon Conservation Score of each motif residue. (C). Motif secondary structure prediction. **β**: Beta-sheet. **α**: alpha-helix. C: Coil.

**Supplementary Figure 3.**
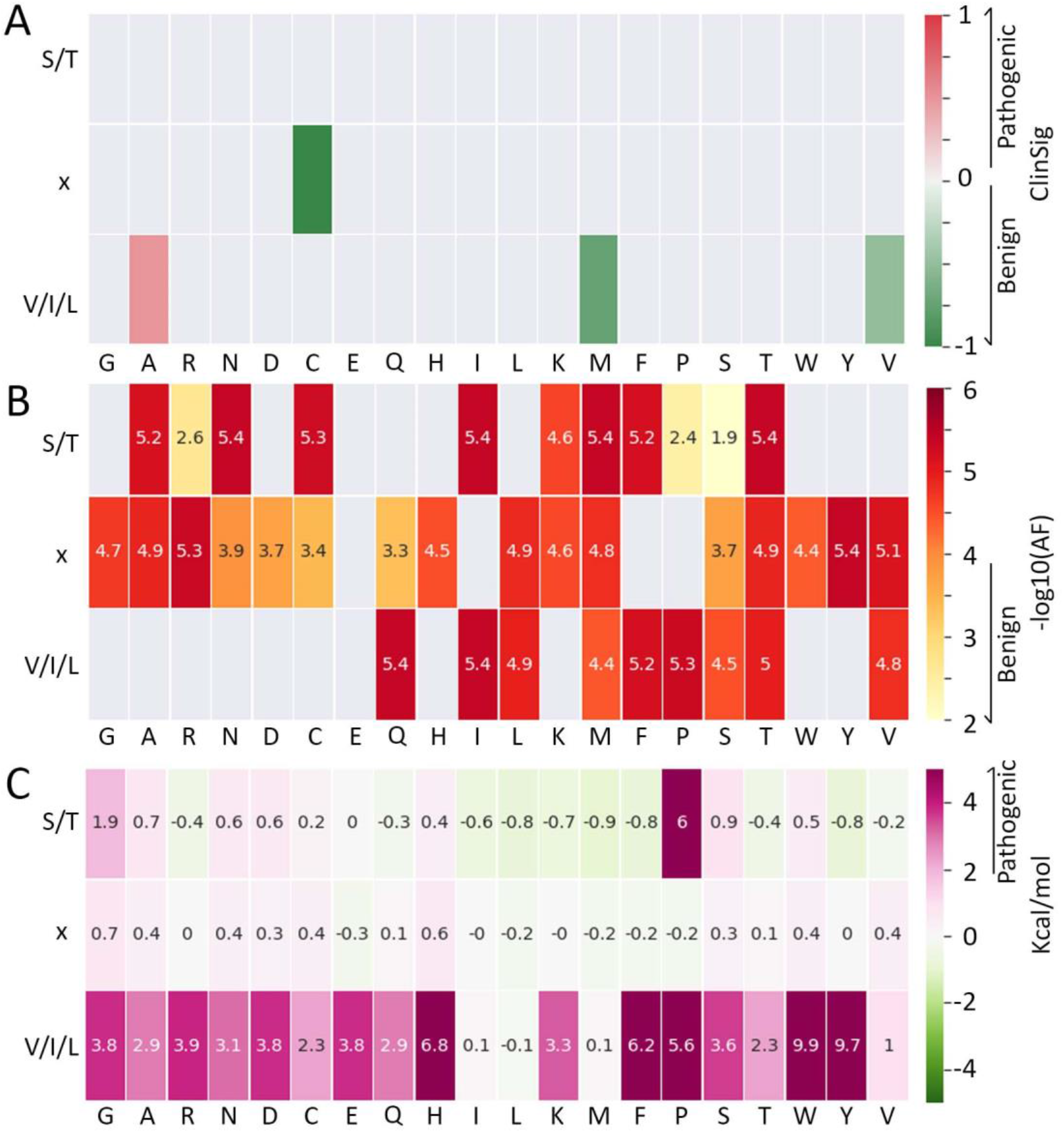
Type 1 PDZ-binding motif P0 SAS Matrices. (A) ClnSigMatrix. Pathogenic variants are presented in red palette and Benign in green palette. (B) AF Matrix. SAS with −log10(AF) < 4 are colored in yellow and were considered as Benign. (C) ΔΔG matrix (in kcal/mol). Values > 2 kcal/mol were considered as pathogenic, while values below 1 kcal/mol were considered as probably tolerated for the interaction.

**Supplementary Figure 4.**
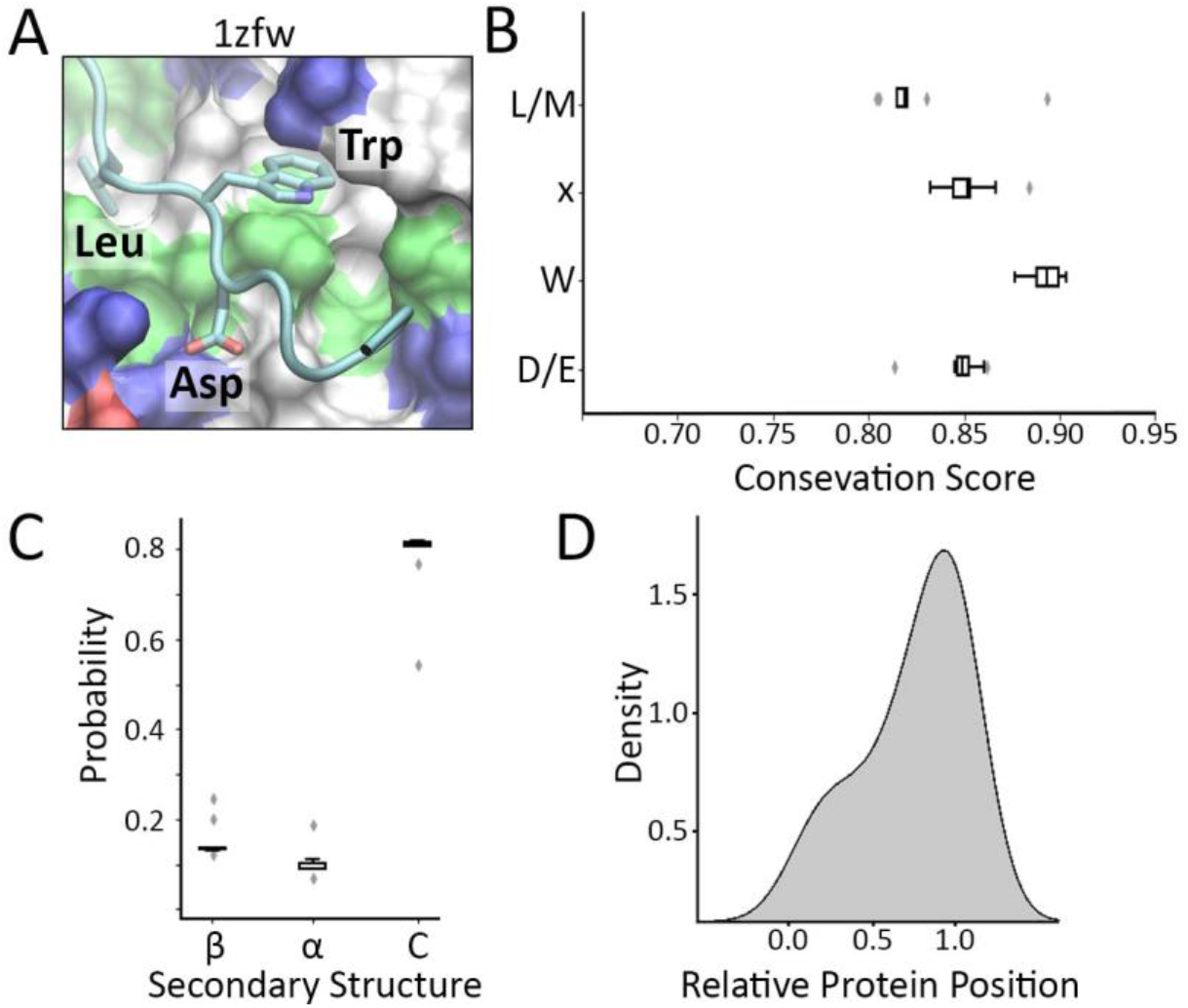
Experimentally tested tryptophan-acidic motifs. (A). Representative crystal structure of the tryptophan-acidic motif and KLC-TPR domains complex. TPR-domains are colored by residue type. (B). Box-plot showing the Jensenn-Shannon Conservation Score of each motif residue. (C). Motif secondary structure prediction. **β**: Beta-sheet. **α**: alpha-helix. C: Coil. (D). Histogram representing the distribution of the relative motif position with respect to the whole protein.

**Supplementary Figure 5.**
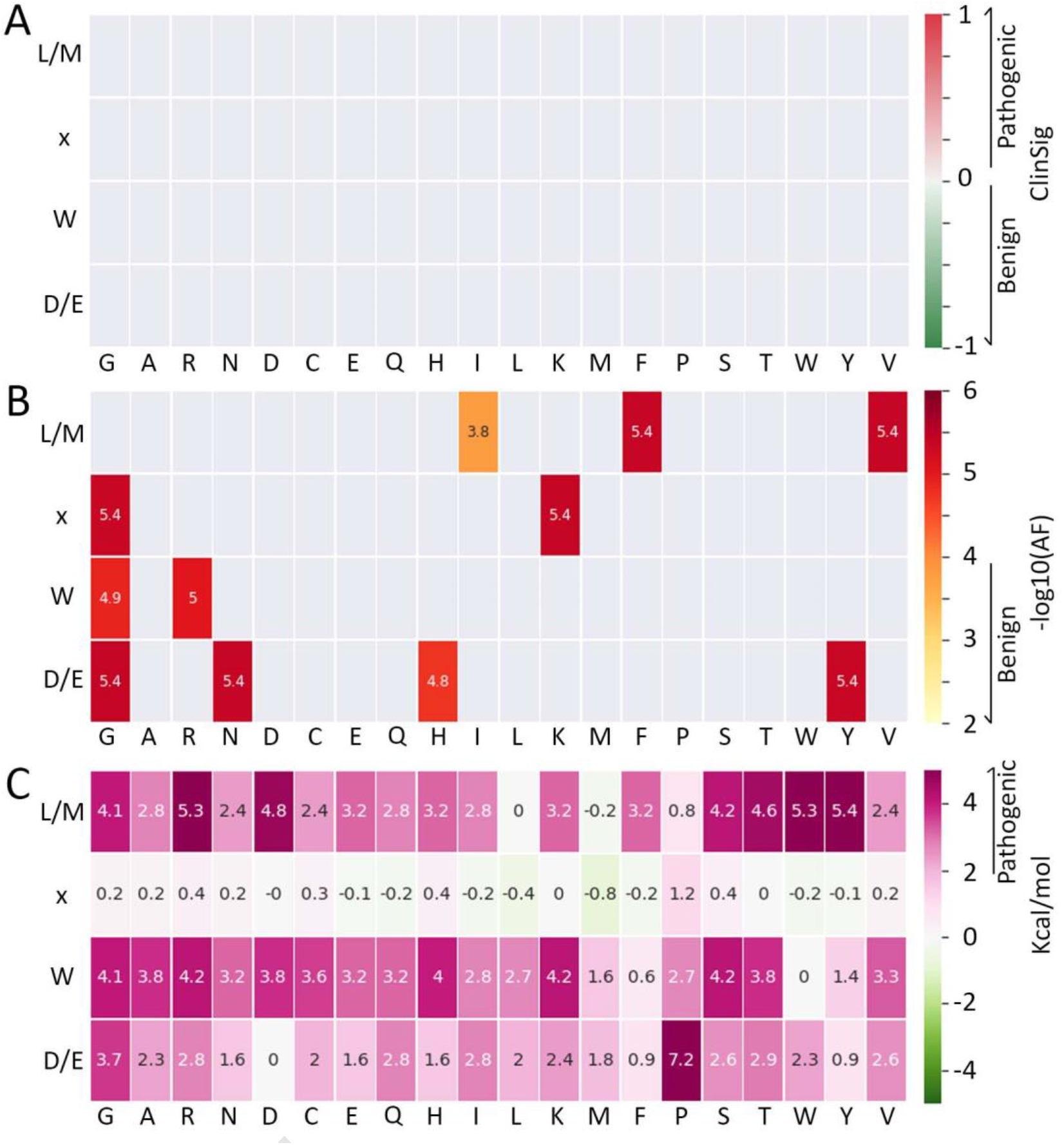
Tryptophan-acidic motif P0 SAS Matrices. (A) ClnSigMatrix. Pathogenic variants are presented in red palette and Benign in green palette. (B) AF Matrix. SAS with −log10(AF) < 4 are colored in yellow and were considered as Benign. (C) ΔΔG matrix (in kcal/mol). Values > 2 kcal/mol were considered as pathogenic, while values below 1 kcal/mol were considered as probably tolerated for the interaction.

**Supplementary Figure 6.**
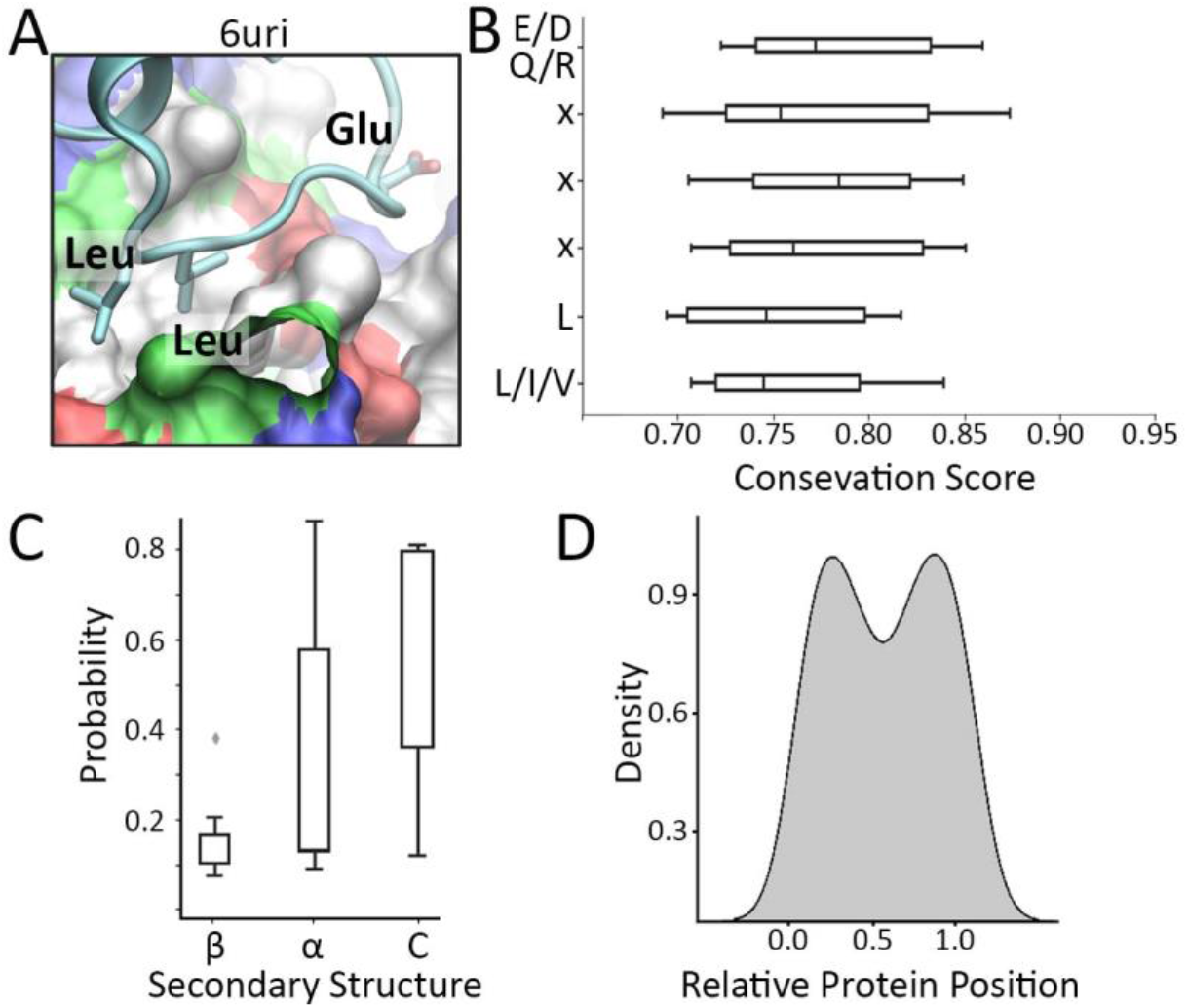
Experimentally tested Acidic di-Leucine motifs. (A). Representative crystal structure of the acidic di-leucine motif and sigma subunit complex. Sigma subunit is colored by residue type. (B). Box-plot showing the Jensenn-Shannon Conservation Score of each motif residue. (C). Motif secondary structure prediction. **β**: Beta-sheet. **α**: alpha-helix. C: Coil. (D). Histogram representing the distribution of the relative motif position with respect to the whole protein.

**Supplementary Figure 7.**
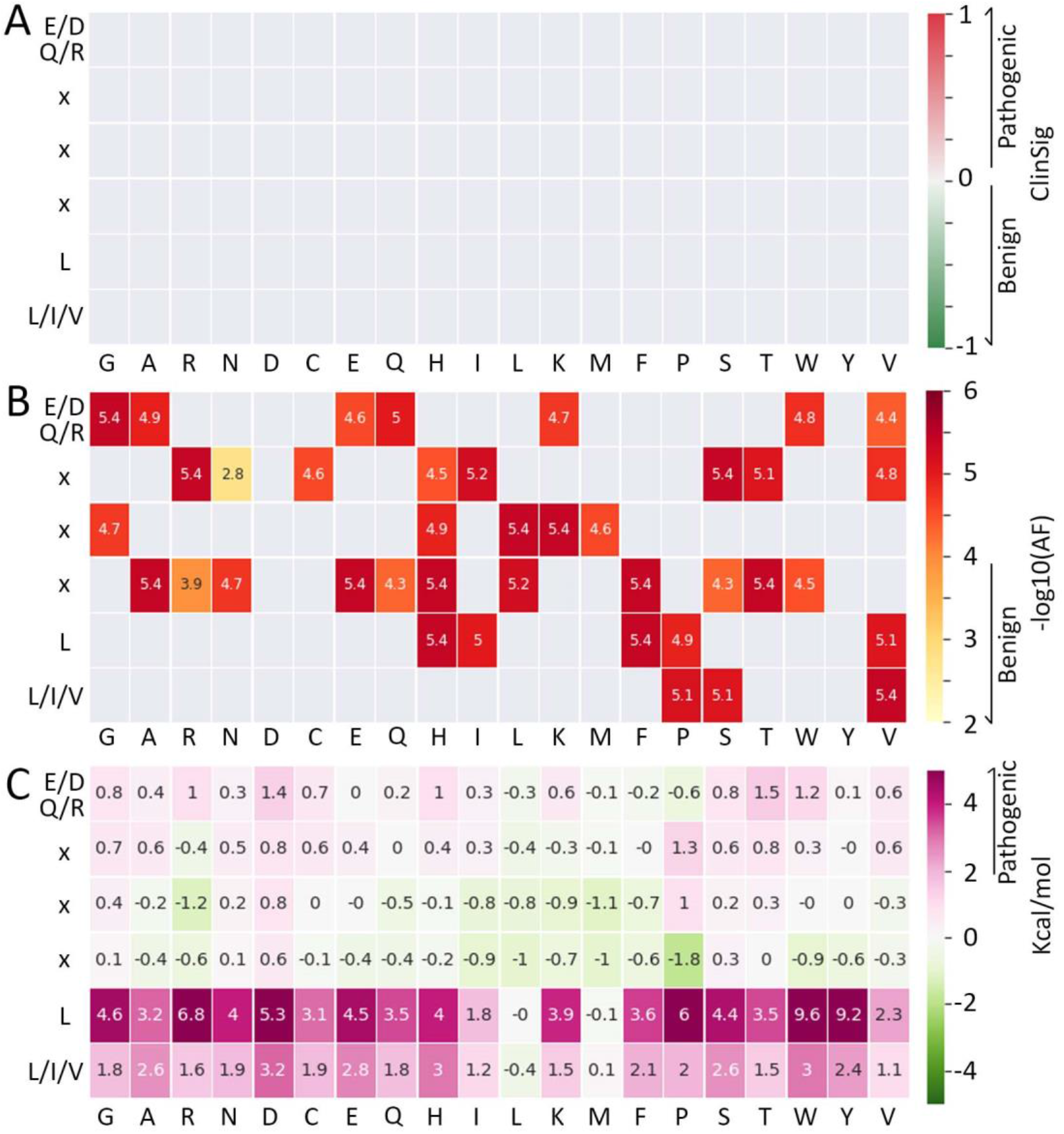
Acidic di-Leucine motif P0 SAS Matrices. (A) ClnSigMatrix. Pathogenic variants are presented in red palette and Benign in green palette. (B) AF Matrix. SAS with −log10(AF) < 4 are colored in yellow and were considered as Benign. (C) ΔΔG matrix (in kcal/mol). Values > 2 kcal/mol were considered as pathogenic, while values below 1 kcal/mol were considered as probably tolerated for the interaction.

**Supplementary Table 1. Discarded and predicted motifs after MotSASi iteration.** UniProtIDs are detailed for each predicted protein. The number after UniProtID represent the order of appearance of the predicted motif in the protein sequence starting from 0 (0 is the first motif in the protein sequence, 1 is the second, etc. For proteins that only have one motif detected is also 0).

